# An epitope-resurfaced virus-like particle can induce broad neutralizing antibody against four serotypes of dengue virus

**DOI:** 10.1101/351700

**Authors:** Wen-Fan Shen, Jedhan Ucat Galula, Jyung-Hurng Liu, Mei-Ying Liao, Chang-Hao Huang, Yu-Chun Wang, Han-Chung Wu, Jian-Jong Liang, Yi-Ling Lin, Matthew T. Whitney, Gwong-Jen J. Chang, Sheng-Ren Chen, Shang-Rung Wu, Day-Yu Chao

## Abstract

Dengue fever is caused by four different serotypes of dengue virus (DENV) which is the leading cause of worldwide arboviral diseases in humans. Virus-like particles (VLPs) containing flavivirus prM/E proteins have been demonstrated to be a potential vaccine candidate; however, the structure of dengue VLP is poorly understood. Herein we show for the first time that mD2VLP particles possess a T=1 icosahedral symmetry with a groove located within the E-protein dimers near the 2-fold vertices that exposed highly overlapping, cryptic neutralizing epitopes through cryo-electron microscopy reconstruction. Mice vaccinated with highly matured virus-like particles derived from DENV serotype 2 (mD2VLP) can generate higher cross reactive (CR) neutralization antibodies (NtAbs) and were protected against all 4 serotypes of DENV through clonal expansion supported by hybridoma and B-cell repertoire analysis. Our results revealed that a “epitope-resurfaced” mature-form dengue VLP has the potential to induce quaternary structure-recognizing broad CR NtAbs.

## Introduction

Dengue virus (DENV), a member of the family *Flaviviridae*, is a mosquito-borne pathogen with four distinct serotypes, DENV-1 to DENV-4(Lindenbach and Rice). It has been estimated that DENV infects about 390 million individuals globally each year, resulting in 96 million clinically apparent infections ranging from mild fever to the life-threatening dengue hemorrhagic fever (DHF) or dengue shock syndrome (DSS)(Bhatt et al., 2013). Although the chimeric yellow fever 17D-derived, tetravalent dengue vaccine (CYD-TDV) has recently been approved by the governments of a few DENV-circulating countries, the unexpected low vaccine efficacy in dengue-naïve children or children younger than 6 years old has limited the use of this vaccine in the 9-45 age group living in endemic countries(Guy and Jackson, 2016). Developing a new strategy by either improving the efficacy of CYD-TDV(Aguiar et al., 2016;Halstead, 2017;World Health Organization, 2017) or the second-generation vaccine candidates(Kirkpatrick et al., 2016) is essential to broaden the coverage for all vulnerable populations.

The envelope (E) protein of DENV on the surface of viral particles is the major target of neutralizing antibodies (NtAbs)(Roehrig, 2003;Pierson et al., 2008). The ectodomain of the E protein contains three distinct domains, EDI, EDII, and EDIII, which are connected by flexible hinges to allow rearrangement of domains during virus assembly, maturation and infection(Modis et al., 2003). However, the immune response from humans who have recovered from primary DENV infections is dominated by cross-reactive (CR), non-NtAbs which recognize mainly pre-membrane (prM) or fusion loop of the E (FLE) protein(Dejnirattisai et al., 2010). During virus replication, newly synthesized DENVs are assembled as immature particles in the endoplasmic reticulum at neutral pH, followed by translocation through the trans-Golgi network and low-pH secretory vesicles(Chambers et al., 1990). The pr portion of the prM protein, positioned to cover the fusion loop (FL) peptide at the distal end of each E protein, prevents premature fusion during the virion maturation process along the egress pathway from the infected cells(Li et al., 2008;Kostyuchenko et al., 2013). For DENV to become fully infectious, the pr molecule has to be removed by a cellular furin protease during the egress process(Rodenhuis-Zybert et al., 2010). However, furin cleavage in the low-pH secretory vesicles is thought to be inefficient; hence, DENV particles released from infected cells are heterogeneous populations with different degrees of cleavage and release of the pr portion of prM to form the mature membrane (M) protein(Pierson and Diamond, 2012). Immature DENV (imDENV) particles consist of uncleaved prM protein and partially immature particles (piDENV) containing both prM and M proteins, while fully mature DENV (mDENV) contains only M protein(Junjhon et al., 2010;Pierson and Diamond, 2012). Functional analyses have revealed that a completely immature flavivirus lacks the ability to infect cells unless in the presence of anti-prM antibodies through a mechanism called antibody-dependent enhancement (ADE) of infection(Dejnirattisai et al., 2010;Rodenhuis-Zybert et al., 2010;Rodenhuis-Zybert et al., 2011;da Silva Voorham et al., 2012). ADE plays an important role in dengue pathogenesis, and is potentially modulated by the antibody concentration and the degree of virion maturity (Nelson et al., 2008;Rodenhuis-Zybert et al., 2015).

Virus-like particles (VLPs) containing flavivirus prM/E proteins have been demonstrated to be a potential vaccine candidate, since their ordered E structures are similar to those on the virion surface and also undergo low-pH-induced rearrangements and membrane fusion similar to viral particles(Allison et al., 1995). Also, VLP vaccines present several advantages since they are highly immunogenic, non-infectious, and accessible to quality control as well as increased production capacity. However, we theorized that the process of VLP maturation might be similar to that of dengue viral particles, with heterogeneous pr-containing structures. In this study, we engineered and characterized the structure of mature DENV-2 VLPs through cryo-EM. The immunological properties of mature VLPs were further compared with its immature counterparts.

## Results

As we theorized, the process of VLP maturation are similar to that of dengue viral particles so that the wild-type particles (wtD2VLP) produced from the cells are partially immature containing both prM and M proteins (Figure S1 in Supplementary Material). To overcome this problem, we engineered the DENV-2 VLPs as completely mature particles by manipulating the furin cleavage site at the junction of pr and M in a DENV-2 VLP-expressing plasmid(Chang et al., 2003). For comparison, we also generated immature VLP (imD2VLPs) by mutating the minimal furin cleavage motif REKR to REST(Li et al., 2008), which resulted in completely uncleaved prM as detected by western blot and ELISA (Figure 1). Generation of mature DENV-2 VLP (mD2VLP) was much more challenging. First, we performed multiple sequence alignments of the pr/M junction from several flaviviruses, including the distance-related cell fusion agent virus (CFAV), and analyzed their furin cleavage potential using the algorithm PiTou 2.0(Tian et al., 2012). DENV-3 pr/M junction had the lowest predicted Pitou score of 6.87, while WNV had the highest score of 15.4 (Figure 1A). Secondly, we determined the cleavage efficiency of D2VLP by replacing P1-8 at the pr/M junction site of DENV-2 with sequences from different flaviviruses. Although the Pitou prediction of the CFAV pr/M junction did not result in a high score for cleavage, replacement with P1-8 of CFAV resulted in the most efficient furin cleavage for D2VLP, with only 31% of prM remaining uncleaved as compared to the 100% uncleaved imD2VLP (Figure 1A). An additional mutation in the P3 residue from E to S of the CFAV P1-8 construct reduced the prM signaling to 13.7% of that of imD2VLP. We thus named this construct with the replacement of CFAV P1-8 plus the P3 E to S mutation as mD2VLP. The pr/M cleavage of this construct was increased to nearly 90% (Figure 1B). Similar to wtD2VLP purified from plasmids transfected culture supernatants, the mD2VLP and imD2VLP showed consistent conformational integrity as ascertained by similar equilibrium banding profiles in 5-25% sucrose density gradients, as well as comparable particle size distributions as determined by negative staining electron microscopy, and complex glycosylation on the E proteins (Figure S2 in Supplementary Material).

**Fig. 1.**
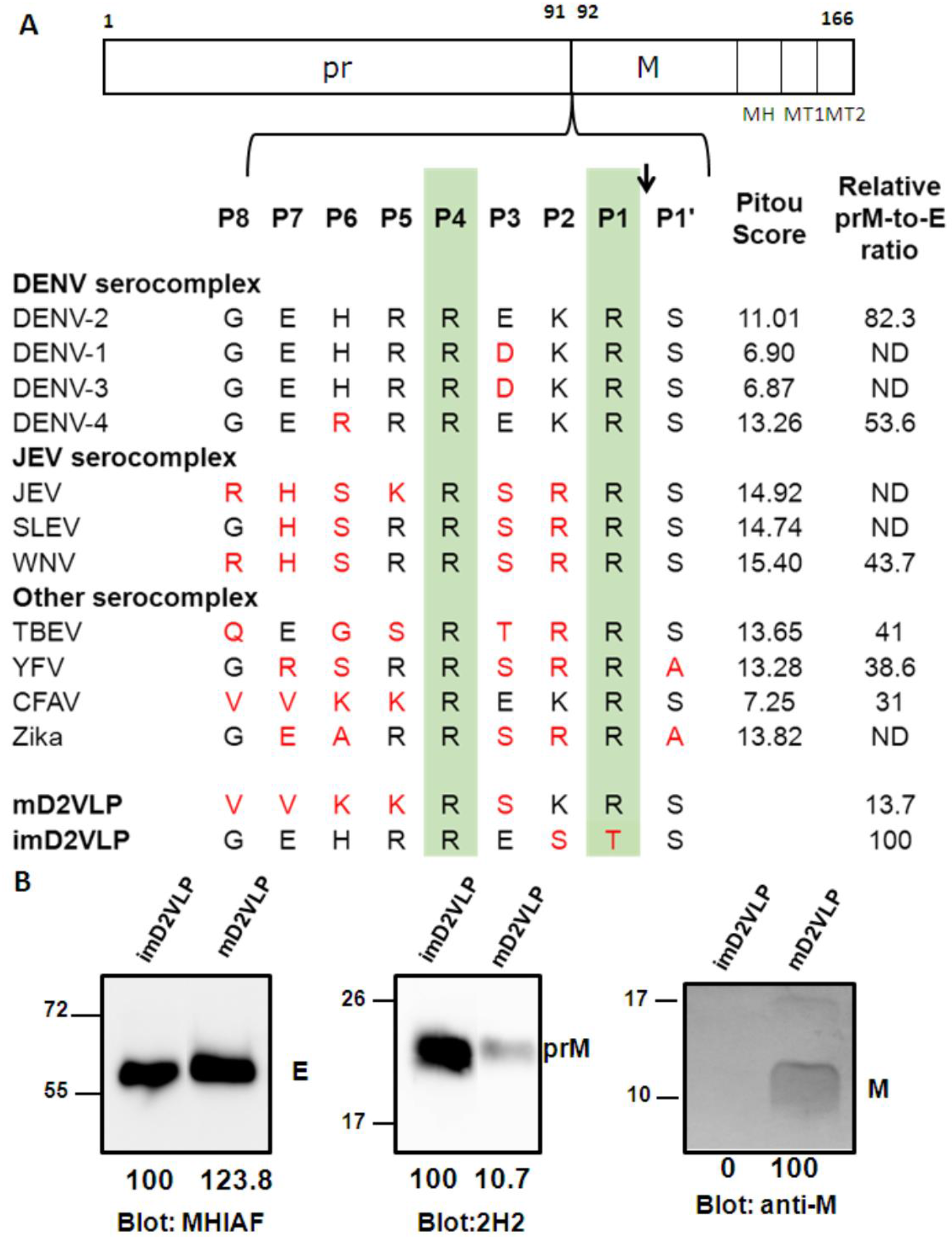
Comparison of the prM junction cleavage efficiency among different DENV-2 virus-like particles (D2VLPs). (A) Schematic drawing of the prM protein. The C-terminal of the prM protein contains an a-helical domain (MH) in the stem region, followed by two transmembrane domains (MT1 and MT2). Numbers refer to the position of the amino acids in the polyprotein starting at the first amino acid of prM according to DENV-2 (NP_056776). Single letter designations of amino acid sequence alignment of representative strains from different serocomplexes of flaviviruses at the prM junction site includes dengue virus serotypes 1-4 (DENV-1 to DENV-4), Japanese encephalitis virus (JEV), St. Louis encephalitis virus (SLEV), West Nile virus (WNV), tick-borne encephalitis virus (TBEV), yellow fever virus (YFV), cell-fusion agent virus (CFAV), Zika virus (ZKV), immature DENV-2 VLP (imD2VLP) and mature DENV-2 VLP (mD2VLP). Numbers with P in the beginning refer to the positions of the amino acid relative to the prM cleavage site in the proximal direction (without apostrophe) and the distal direction (with apostrophe). Protein sequences were aligned, with the key P1 and P4 positions within the furin cleavage sites highlighted. The arrow indicates the prM cleavage site. The amino acids in red indicate the residues different from DENV-2. The PiTou 2.0 furin cleavage prediction scores are shown on the right for each sequence and the higher scores indicate the higher efficiency of furin cleavage. The relative quantity of prM and E of the wild-type and mutant DENV-2 VLP with P1-8 replacement (from other flaviviruses as shown) was measured by ELISA using MAb 3H5 (specific to E domain III of DENV-2) and MAb 155-49 (specific to DENV prM). The relative prM-to-E ratios were calculated by absorbance for prM/absorbance for E protein with reference to imD2VLP, whose pr portion was set as 100% uncleaved, as shown on the right. ND: not determined. Data are presented as means from three representative ELISA experiments with two replicates. (B) Culture supernatants of mD2VLP and imD2VLPs were collected and purified after electroporation with the respective plasmids. Five micrograms of proteins were loaded onto a 12% non-reducing Tricine-SDS-PAGE. E, prM and M proteins were assayed by Western blot using mouse hyper-immune ascitic fluids (MHIAF, 1:2000), MAb 2H2 (0.5 μg/mL) and anti-M protein mouse sera (1:25), respectively. E and prM proteins were visualized with enhanced chemiluminesence (ECL); however, M protein was visualized by TMB substrate to avoid high background. The number below each blot shows the relative densitometric quantification of E, prM and M protein bands by Bio-D1D software.

Previous study has shown molecular organization of recombinant VLPs from tick-borne encephalitis virus (TBEV)(Ferlenghi et al., 2001). However, the cryo-EM results showed low resolution due to particle size heterogeneity and no immunological data can be derived from such structure. To better understand if the structural organization of mD2VLP is similar to that of TBEV and its immunological characteristics, purified mD2VLPs were subjected to cryo-EM analysis. The results demonstrated that the mD2VLPs had a variable size distribution (Figure S3 in Supplementary Material). The size variation made improving the resolution challenging, however we were able to determine the overall morphology and E arrangement of mD2VLP based on current achieved resolution. Using a dengue virus-specific Mab (MAb32-6)(Li et al., 2012), immunogold labeling detected the mD2VLPs as spherical particles with a diameter of ~31 nm (Figure S4 in Supplementary Material), which was also the major population as shown in Supplementary Figure S3B. Besides, the particles with diameter of ~31nm had more solid features in 2D image analyses (Figure S3 in Supplementary Material) and were able to obtain stable structure than other groups. Therefore, we continued and focused on this population for further 3D reconstruction and image analyses. The results showed that mD2VLPs with the diameter of ~31nm at a resolution of 13.1Å had a hollow structure and smooth surface with protrusions around the 5-fold positions (Figure 2A, left, and Figure S5 in Supplementary Material). Fitting the atomic E and M surface proteins (PDB: 3J27) into the cryo-EM density map showed that the mD2VLPs have 60 copies of E packed as dimers in a T=1 icosahedral surface lattice (Figure 2A, right). The apparent differences of structural features on the mD2VLPs compared to the mature native virion particles(Kostyuchenko et al., 2013;Zhang et al., 2013a) was a slimmer lipid bilayer and the rearrangements of the surface proteins. First, the central section through the viron and m2DVLP reconstructions showed that the bilayer of m2DVLP was relatively thinner than that of viron. The distance between the exterior leaflet and the interior leaflet of m2DVLP and virion was 12Å and 14Å, respectively (Figure 2B). A similar T=1 icosahedral symmetry of the E-protein arrangement and thinner lipid bilayer were also found in TBEV VLPs(Ferlenghi et al., 2001). Second, interpretation of the map showed that E dimer subunits moved apart from each other causing less density in the intra-dimeric interphase (Figure 3C). Because of this loose interaction, a groove located within the E-protein dimer near the icosahedral 2-fold vertices on mD2VLP was noted (Figure 3A, right, and Figure S6 in Supplementary Material). Third, the central section showed that there was a solid density in the H-layer, which is composed of H helices of E and M protein stem regions, while the density in the T-layer, which contains transmembrane helices of E and M proteins, was weaker in the VLP (Figure 2B). Each E protein consists of an E ecto-domain and a stem region that connects the ecto-domain to its transmembrane region. The stem contains two α-helices, one of which interacts with the viral lipid membrane and the other with the E ecto-domain. This looser interaction of E protein dimers was further stabilized by the E protein rearrangement and the closer interaction with one of the α-helices, which resulted in the solid density beneath the ecto-domain as shown in Figure 2B. It is worth noting that the stem region of this mD2VLP was replaced into Japanese Encephalitis viral (JEV) sequence; it was previously suggested that this replacement increased the hydrophobicity for the interaction with the lipid membrane and enhanced the secretion of dengue VLP(Purdy and Chang, 2005).

**Figure 2.**
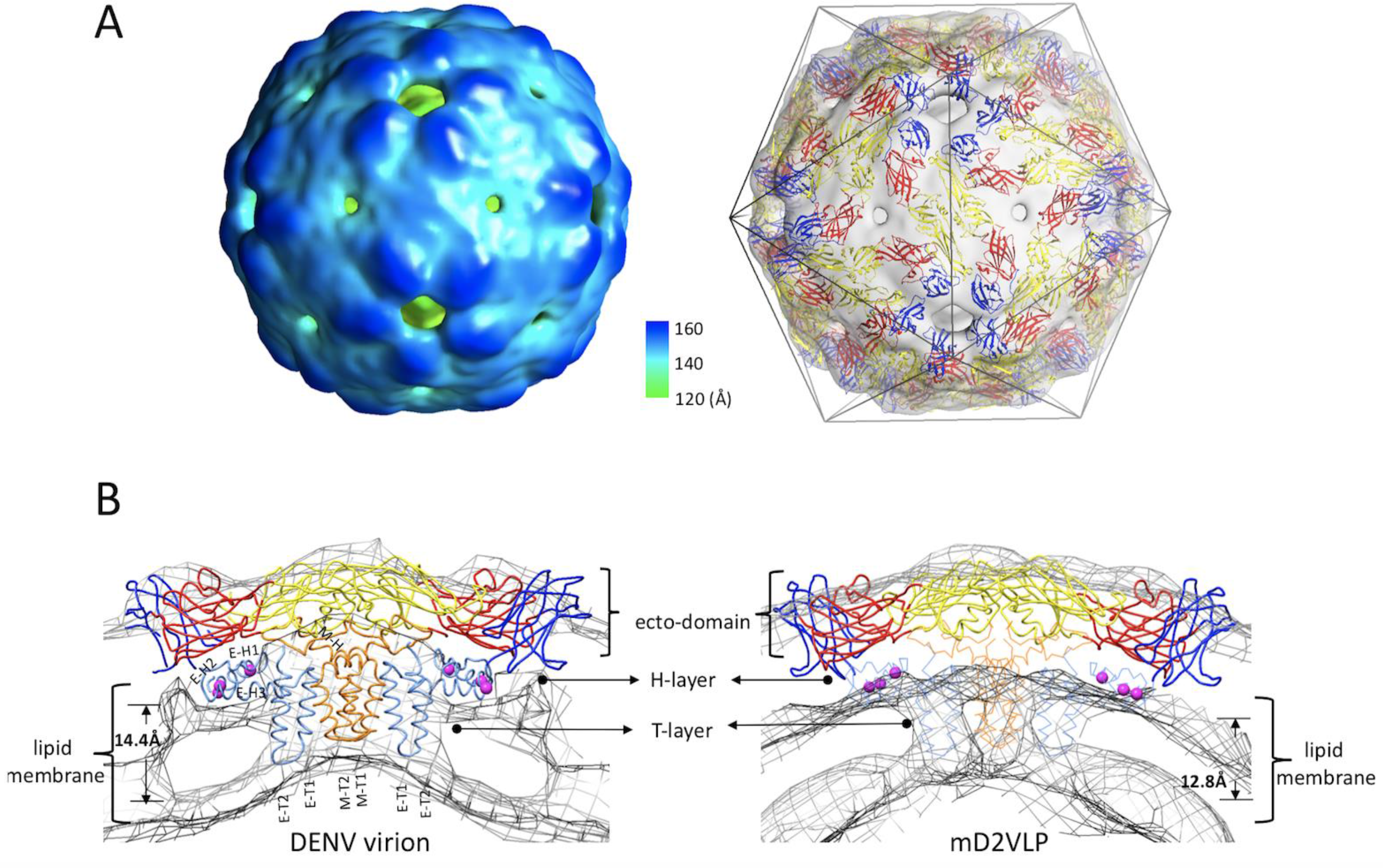
The structure of mature form virus-like particles of dengue virus serotype 2 (mD2VLP). **(A)** The reconstructed cryo-EM map of the DENV VLPs (left panel) were presented with the radial color-code indicated (left), the size of the particle is 31nm. Fitting of atomic E, M surface protein (PDB: 3J27) into cryo-EM density map (right) showed that the VLP has 60 copies of E in a T=1 arrangement. The density map was shown as a transparent volume rendering into which was fitted the backbone structures of the E ecto-domain. The domains I, II, and III were highlighted in red, yellow and blue, respectively. The cage indicated the icosahedral symmetry. **(B)** The cross section showed the fitted E:M:M:E heterotetramer (PDB: 3J27) into DENV virion (left) and into mD2VLP (right). The map density was in mesh presentation. The atomic model of E:M:M:E heterotetramer was showed in ribbon. Domain definition of dengue E was the same as the previous description, the transmembrane domain of E was colored as light blue, the M protein was in orange color. It was clear that the density of H-layer which is composed of E-H1 to E-H3 and M-H was more solid while the density in T-layer which contains E-T1, E-T2, M-T1 and M-T2 was weak in VLP than in virion. The residues at 398, 401 and 412 in E-H1 of JEV sequence which were proved to play important role in promoting extracellular secretion^28^ were shown as magenta spheres. The transmembrane, perimembrane helices and lipid bilayer were labelled, the critical measurements were also shown.

Next, we speculated that the structure of mD2VLP with a groove located within the E-protein dimer could affect the epitope presentation on the surface of the particles. First, we calculated the solvent accessibility of mD2VLP and compared the results with that of DENV-2 virion. This rearrangement of E protein under T=1 symmetry on the VLP surface exposed more accessible epitopes (48.2%; 191 amino acid with ≥30% solvent accessibility among 396 amino acids of the entire E protein) compared to DENV-2 virions (43.4%; 172 amino acid), particularly located at fusion loop, aa 239, aa 251-262 of EDII which together formed the groove, and at A strand, cd loop and G strand of EDIII which surrounded a 5-fold axis (Figure 3A, left, Figure S7 in Supplementary Material). Footprint analysis (Figure 3B) showed that those cryptic residues on the E protein, which were previously buried inside of the native DENV virion and only interacted with the broad NtAbs while the virion “breathed” (Lok, 2016), were better exposed on the m2DVLP. Specifically, the epitopes interacting with MAb 1A1D-2, which binds to the virus at 37°C(Lok et al., 2008); MAb 2D22, which could block two-thirds of all dimers on the virus surface, depending on the strain(Fibriansah et al., 2015a); and the ‘E-dimer-dependent epitope’, which is recognized by broadly neutralizing MAbs EDE1 and 2(Rouvinski et al., 2015), were all well-exposed in VLPs without steric hindrance. Second, a panel of murine MAb, recognizing different domains of E protein and used previously for antigenic mapping(Crill et al., 2012), was subjected to binding-ELISA (Figure S8 in Supplementary Material). The results confirmed that most of the Mabs preferentially binds to mD2VLP, in particularly Mabs recognizing domain I/II (Figure 4).

**Figure 3.**
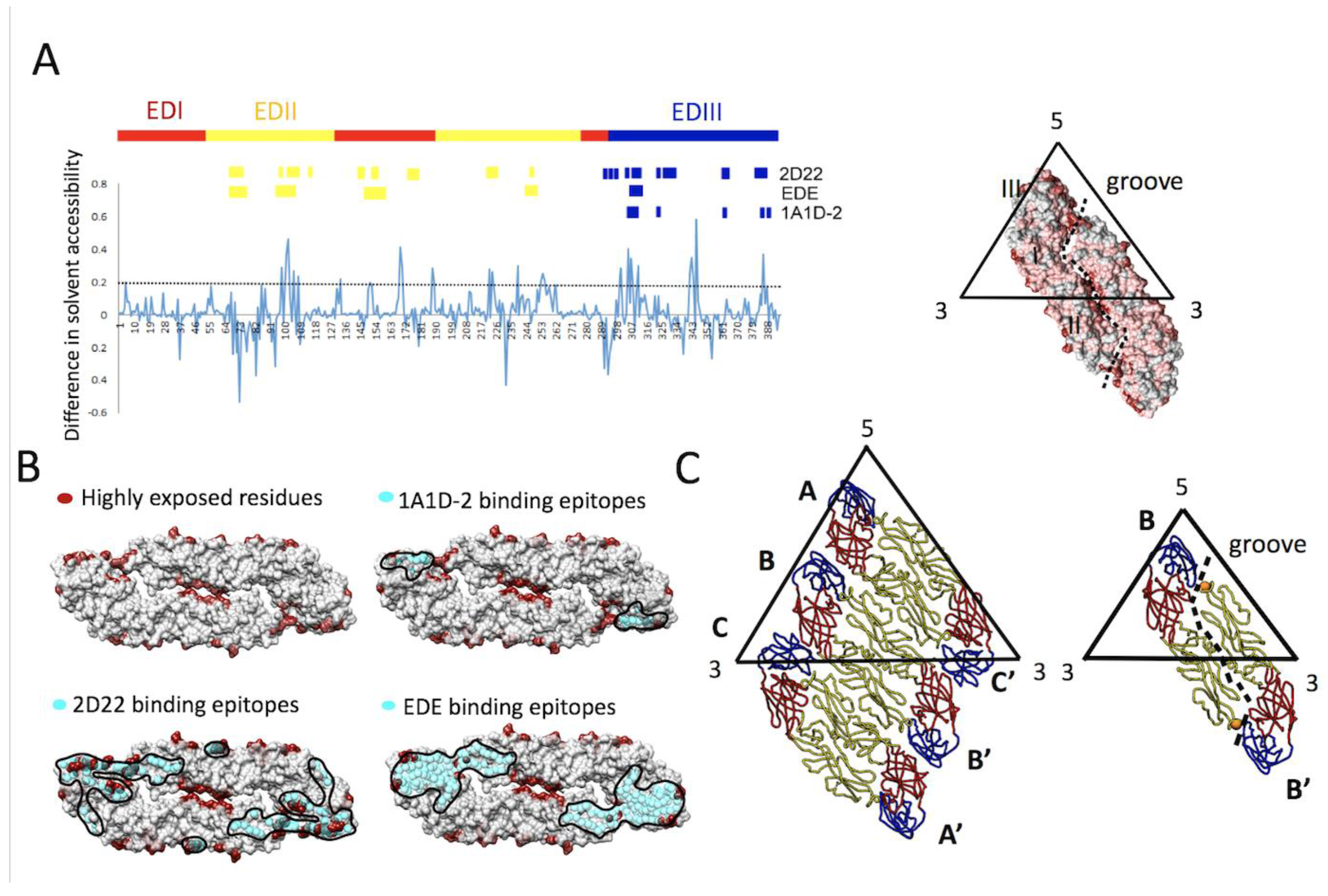
Solvent accessibility of dengue virus serotype 2 virus soluble envelope (sE) protein (A, left) The plot of difference in relative solvent-accessible surface area (Δ%SASA). A positive value of %SASA meant the residue became more exposed when the particle assembly shifts from virion (3 copies of E dimers per icosahedral asymmetric unit) to VLP (1 copy of E dimers per icosahedral asymmetric unit). The black dash line indicated the Δ%SASA ≥0.2 which was defined as highly exposed residues in VLP comparing to virion. The high positive values which were focused in the peptide regions such as the fusion loop peptide (including amino acid (aa) residues ranging from 100-110), aa 169-170 at domain I, aa 222-226, aa 239 and aa 251-262 at domain II as well as A strand of domain III (aa 300-308), the cd loop of domain III (aa 342-348) and G strand (aa 386-388) around the 5-fold openings. The residues interacting with MAb 1A1D-2^30^, including residues 305-312, 352, 364, 388 and 390; the residues interacting with MAb 2D22^31^, including residues 67-72, 99, 101-104, 113, 177-180, 225-227, 247, 328, 384-386 (Heavy chain); 148-149, 153-155, 291-293, 295, 298, 299, 307, 309-310, 325, 327, 362-363 (light chain) and the residues interacting with human MAb EDE antibodies^19^, including aa residues 67-74, 97-106, 148-159, 246-249 and 307-314 were indicated. **(A, right)** The high positive peaks (Δ%SASA ≥0.2), low positive peaks (Δ%SASA between 0 and 0.2) and negative peaks (Δ%SASA ≤0) in the plot were colored by dark red, deem red and grey in the E dimer surface rendering. The groove located within E-dimer interface was outlined. **(B)** The highly exposed residues (Δ%SASA≥0.2) which were colored by dark red were shown in the surface rendered E-dimer. The residues in E interacting with MAb 1A1D-2, 2D22 and EDE were in cyan spheres showing that they were highly exposed on the m2DVLP surface. Importantly, the binding footprints of the three antibodies were highly overlapping with footprint of highly exposed residues in m2DVLP, and formed a neutralization sensitive patch on m2DVLPs. The areas of the interacting epitopes are circled by black lines. **(C)** The E protein forming the rafts in virion were shown in the left panel where the three individual E proteins in the asymmetric unit are labeled A, B, and C of the E proteins, in the neighboring asymmetric unit are labeled A’, B’, and C’. The icosahedral 2-fold E protein dimers (B and B’) in m2DVLP have moved apart from each other causing the groove (right). The aa 101 which was responsible for DM25-3 antibody binding were shown in orange spheres.

**Fig 4.**
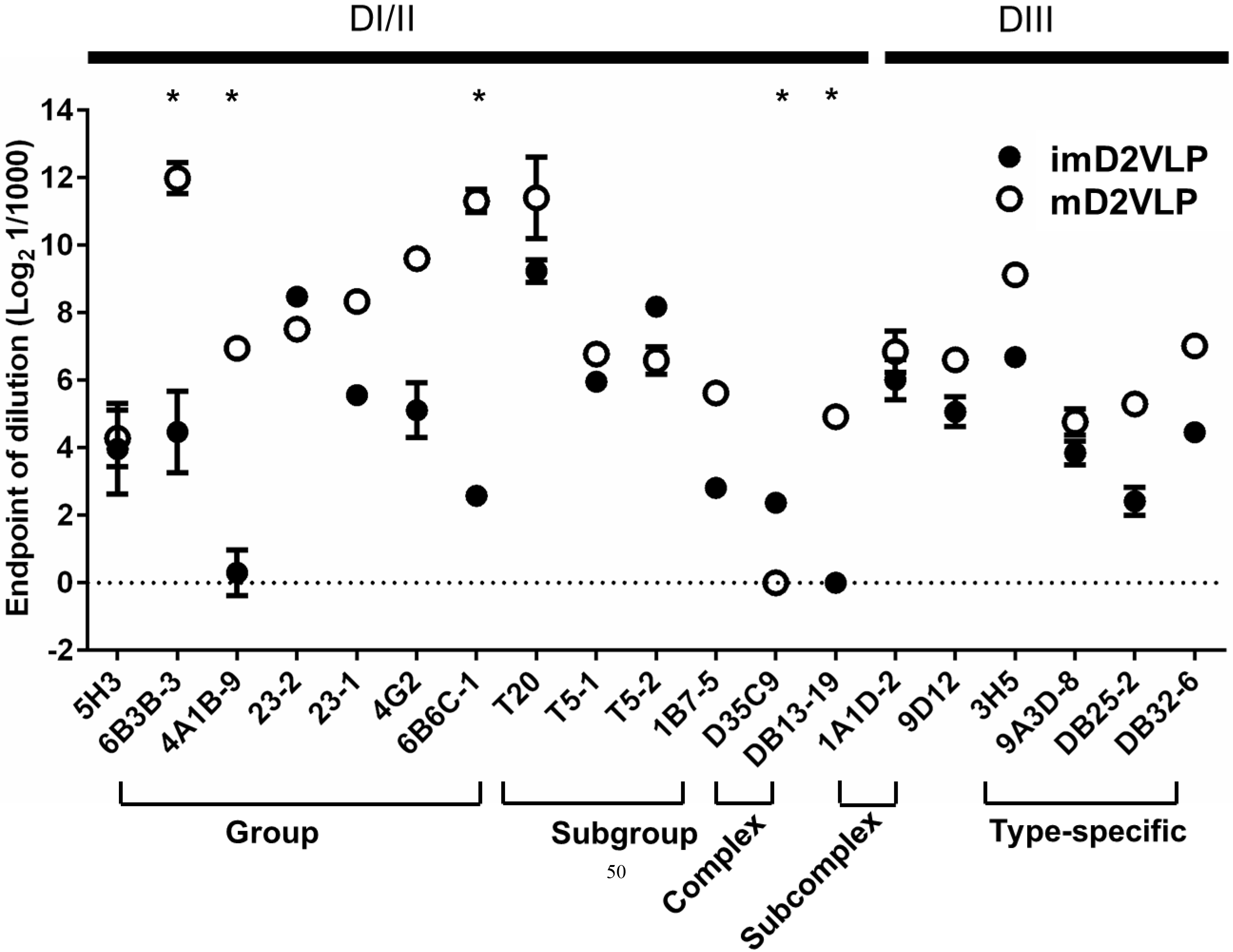
Summary of binding activity of a panel of anti-E monoclonal antibodies (Mabs) to mD2VLP and imD2VLP. The binding activity of each Mab to two types D2VLP was expressed as means of endpoint dilution with standard deviations from duplicates of representative experiments of three. The dash line represents the minimum dilution of Mab used in the experiment. The asterisk symbol represents statistical significance between mD2VLP and imD2VLP in antibody dilution reaching endpoint with p-value <0.05

To further determine the influence of prM cleavage on the immunogenicity of D2VLP, we immunized groups of 4-week-old BALB/c mice with the purified wtD2VLP, imD2VLP or mD2VLP (4 μg/mouse) twice at a 4-week interval. To better represent the neutralizing antibodies at the late convalescent phase, serum samples collected at 8 weeks after the boost were analyzed by antigen-specific ELISA for antibody response against homologous D2VLP antigens with different maturity profiles. In order to precisely quantify the amount of dengue-specific antibodies, we used the same quantity of purified VLPs in the antigen-capture ELISA (Figure 5A). WtD2VLP and imD2VLP induced similar ELISA titers of DENV-2-specific IgG antibody against three VLP antigens. However, stronger reactivity against the mature antigen was noted with mD2VLP immunization (1:15,347 vs. 1:6,381 vs. 2,529 for mD2VLP, imD2VLP and wtD2VLP, respectively) (Figure 5B). We also analyzed the neutralizing ability of these sera against the four serotypes of DENV using a 50% antigen focus-reduction micro neutralization test (FRμNT_50_). The mD2VLP immunization group induced a higher and broader neutralizing antibody response against all 4 serotypes of DENV (FRμNT_50_ for DENV-1=1:331, DENV-2=1:597, DENV-3=1:70 and DENV-4=1:141), as compared to the imD2VLP vaccinated group (FRμNT50 for DENV-1=1:100, DENV-2=1:207 DENV-3=1:64 and DENV-4=1:76) (Figure 5C).

**Figure 5.**
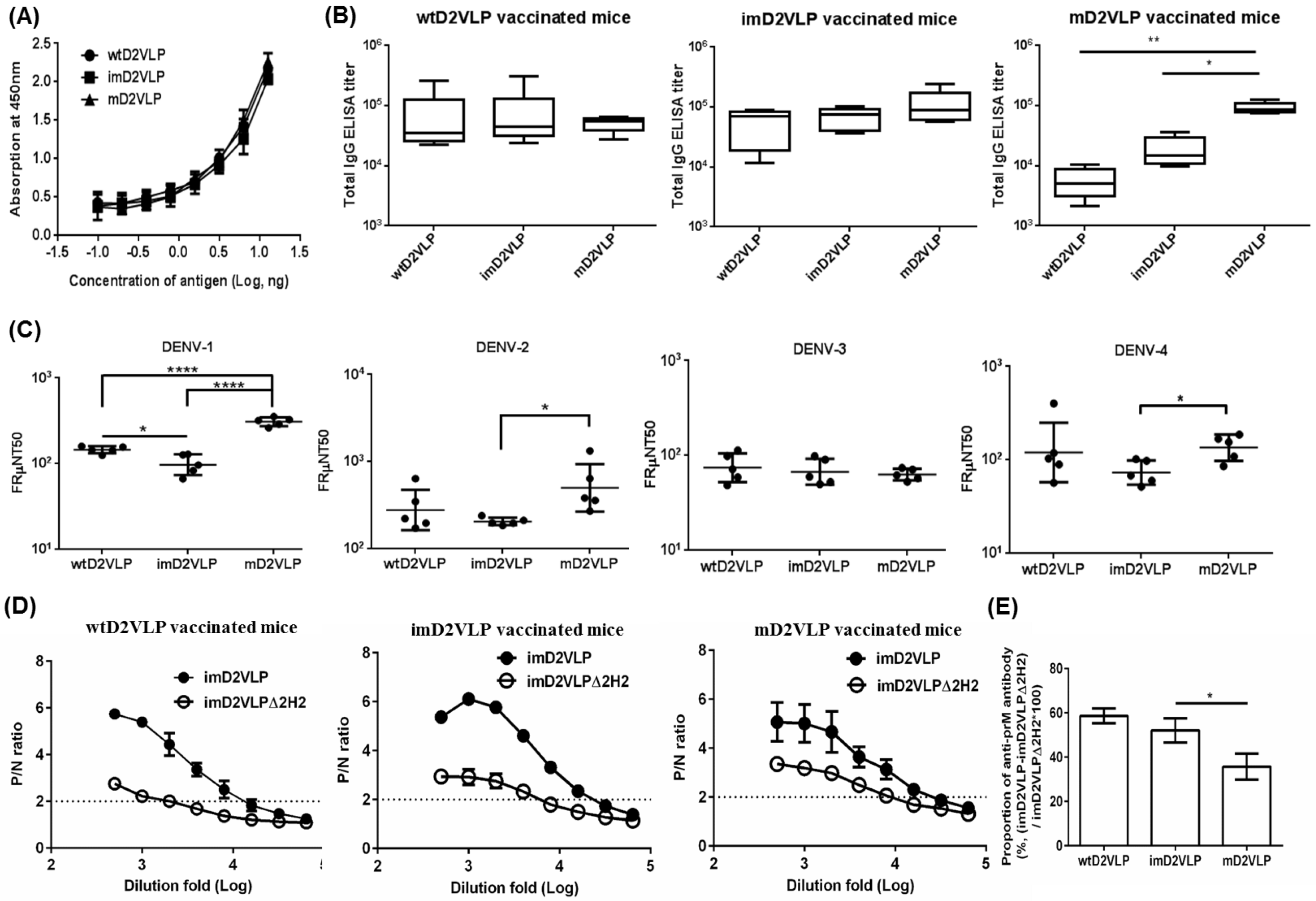
Total antigen-specific IgG, neutralizing titers and proportion of anti-prM antibodies compared among three groups of mice immunized with wtD2VLP, imD2VLP or mD2VLP. **(A)** D2VLPs were concentrated and purified from clarified supernatants. The total protein concentration of purified imD2VLP and mD2VLP were first determined by the Bradford assay and then subjected to antigen-capture ELISA using 2-fold serial dilutions. The standard curve was used to titrate both antigens as equal amounts for the subsequent assays. **(B)** The endpoint IgG titer of 12-week post-immunization mouse sera was measured by antigen-capture ELISA, using equal amounts of homologous and heterologous purified D2VLP antigens. All endpoint titers were log_10_ transformed and depicted as geometric means with 95% confidence intervals. **(C)** The neutralizing antibody titers at 50% antigen focus-reduction micro neutralization (FRμNT50) in Vero cells infected with DENV-1 to 4. **(D)** Binding reactivity of serial dilutions of anti-wtD2VLP (left), anti-imD2VLP (center), and anti-mD2VLP (right) mouse sera were analyzed by ELISA using equal amounts of wild-type (imD2VLP) and mutant imD2VLP (Δ2H2) antigens. **(E)** Proportions of 2H2-like, anti-prM antibodies from two different D2VLP immunization groups were calculated based on the formula 100*[(OD450imD2VLP-0D450A2H2)/0D450imD2VLP] at a 1:1000 dilution of mouse sera. All data presented are based on a representative of three independent experiments with two replicates from n=5 mice sera per group per experiment and expressed as mean ± SEM. The statistical significance was determined using the two-tailed Mann-Whitney *U* test to account for non-normality of the transformed data. *, *P* <0.05; **, *P* <0.01; ****, *P* <0.0001.

Since the amino acid sequence is identical except for the mutations at the furin cleavage site, the difference in antibody binding and neutralizing activity between mD2VLP and imD2VLP vaccinated mice sera could result from the difference in induction of CR anti-prM non-NtAbs or anti-E NtAbs recognizing structure-dependent epitopes. To address whether the higher neutralization activity induced by mD2VLP was partly due to the reduction of anti-prM antibodies that are known to be cross-reactive but have no neutralizing activity(Dejnirattisai et al., 2010), we performed epitope-blocking ELISAs by using an anti-prM-specific monoclonal antibody (MAb 2H2). MAb 2H2 blocked only 7.59% of the activity of anti-mD2VLP sera but blocked up to 35.26% of antibodies from the imD2VLP-vaccinated groups (Figure S9 in Supplementary Material). To avoid steric hindrance due to Mab 2H2 binding, we performed site-directed mutagenesis on three amino acids of pr protein based on the following criteria: (1) the conservation of amino acids among all four serotypes; (2) residues interacting with MAb 2H2(Wang et al., 2013); (3) residues not interfering with prM and E interaction(Li et al., 2008). As shown in supplementary Figure S10, K26P was a key residue, which significantly decreased the binding of both MAb 2H2 and 155-49 (another cross-reactive murine Mab recognizing pr protein(Huang et al., 2006)). However, only the mutant A2H2-imD2VLP with F1A, K21D and K26P amino acid triple mutations completely prevent the binding of both MAb 2H2 and 155-49 but not that of DENV-2, E-specific Nt-MAb 3H5 (Figure S11 in Supplementary Material). The sera from immunized mice were tested for their binding ability to the wild-type and mutant-A2H2 imD2VLP. The differences in binding of imD2VLP and A2H2 imD2VLP was greater for the anti-imD2VLP sera than the anti-mD2VLP sera (Figure 5D and E), suggesting that compared to mD2VLP, imD2VLP was more likely to induce 2H2-like anti-prM antibodies. Thus, mD2VLP has the advantage of eliciting lower levels of anti-prM CR antibody and inducing antibodies with greater neutralizing activity.

The preservation of neutralizing epitopes on the surface of VLPs is critical for efficient production of broadly neutralizing antibody responses. Recent studies have suggested that human monoclonal antibodies (MAbs) reactive with all dengue serotypes can neutralize DENV in the low picomolar range. These MAbs have a preference to bind at the envelope dimer epitopes preserved on virion particles with a high degree of maturity(Dejnirattisai et al., 2015;Rouvinski et al., 2015). We next investigated if a broad neutralization response to mD2VLP immunization was due to the induction of antibodies with a preference to bind at the E dimer epitopes preserved on the surface of mature VLP(Dejnirattisai et al., 2015;Rouvinski et al., 2015). The splenocytes from mD2VLP-vaccinated mice were used to perform fusion and generate hybridomas. Among 2,836 hybridomas screened, two MAbs with high reactivities to VLP measured by ELISA and FRμNT_50_ were shown in Figure 6. As shown in Figure 6A, DM8-6 is a serotype-specific MAb reactive only with DENV-2; while MAb DM25-3 recognized all four serotypes of DENV. Next, we tested the neutralizing activity of these two MAbs against four serotypes of DENV. DM8-6 showed good neutralizing activity against DENV-2 at a concentration of 0.037 μg/mL in FRμNT_50_ but poorly neutralized the other three serotypes. Consistent with the ELISA results, DM25-3 neutralized all four serotypes in FRμNT_50_ at 0.32, 0.38, 0.24 and 0.58 μg/mL for DENV-1 to DENV-4, respectively (Figure 6B).

**Figure 6.**
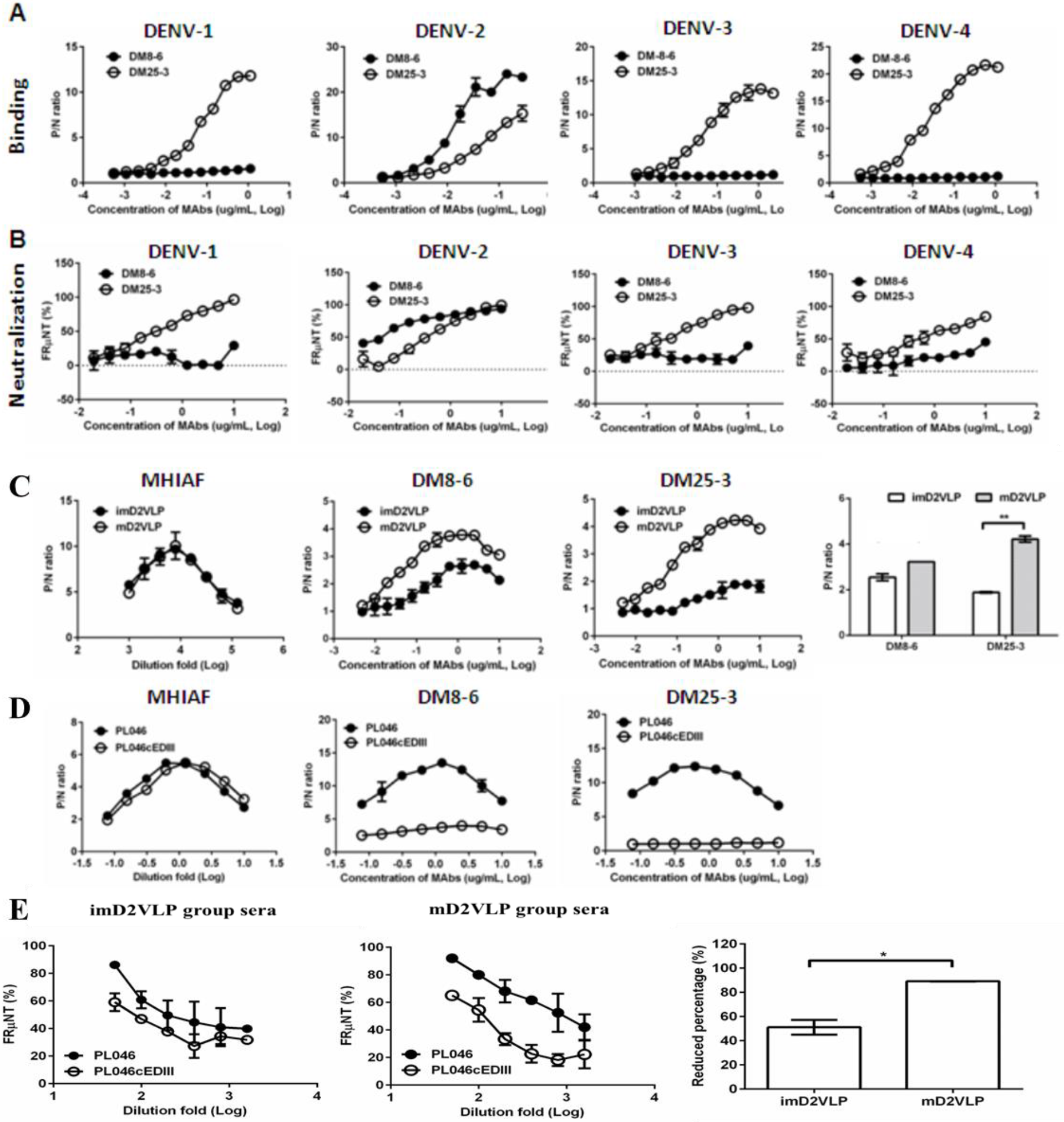
Characterization of murine monoclonal antibodies (MAbs) generated from mouse splenocytes following immunization with mD2VLP. Binding **(A)** and neutralizing **(B)** activities of MAbs DM8-6 and DM25-3 against DENV-1 to 4 were measured by ELISA and the focus-reduction micro-neutralization test (FRμNT). **(C)** Binding curves of MAb DM8-6 (center left) and DM25-3 (center right) against imD2VLP and mD2VLP were performed using ELISAs. Equal amount of both imD2VLP and mD2VLP was properly titrated by antigen-capture ELISA using mouse hyper-immune ascitic fluid (MHIAF) against DENV-2 (left). The difference in binding activities of both MAbs is presented by a bar graph (right). **(D)** A recombinant DENV-2 virus was produced by replacing domain III with a consensus sequence of domain III (PL046cEDIII)^31^ and the binding activity of DM8-6 (center) and DM25-3 (right) was compared with that of parental DENV-2 strain PL046. Equal amount of both PL046 and PL046cEDIII was properly titrated by antigen-capture ELISA using mouse hyper-immune ascitic fluid (MHIAF) against DENV-2 (left). **(E)** FRμNT of two-fold diluted mice sera immunized with mD2VLP and imD2VLP against parental PL046 and PL046cEDIII DENV-2 viruses (n=5 per group per experiment). The differences in FRμNT50 from mD2VLP and imD2VLP immunization groups were converted to bar chart at 1:1000 fold dilution of mice sera. The conversion was based on the formula 100*[ FRμNT50 of (PL046-PL046cEDIII)/FRμNT50 of PL046]. P/N ratio refer to the antibody binding magnitude between designated VLP-containing (P) and VLP-free culture supernatant (N) by dividing the absorbance of P by that of N. The data are presented as means ± SEM from three independent experiments with two replicates. The two-tailed Mann-Whitney *U* test was used to test statistical significance. *, *P* <0.05. **, *P* <0.01.

To determine if MAb DM25-3 recognized quaternary structure-dependent epitopes presented only on mature virion particles(Fibriansah et al., 2015a;Rouvinski et al., 2015), DM25-3 was tested for its binding activity to both mD2VLP and imD2VLP using antigen-capture ELISA. MAb DM25-3 recognized mD2VLP very well, but reacted poorly with imD2VLP (Figure 6C). Next, we performed site-directed mutagenesis on the fusion loop peptide in domain II and A-strand in domain III, both of which are important binding regions for CR group/complex neutralizing antibodies(de Alwis et al., 2011; Tsai et al., 2013) (Figure S12 in Supplementary Material). The results suggested that amino acid E-101 was likely important in the binding site of DM25-3. Residue 101 on the fusion loop of E protein is conserved among all four serotypes of DENV and can only be exposed when virion particles undergo low-pH induced conformational change or during the “breathing” state(Zhang et al., 2013b;Zhang et al., 2015). Figure 3C shows that residue 101 is located at the touching point between EDII and EDIII of the dimeric molecule. Since residue 101 is a critical residue in stabilizing E-protein dimers on DENV virion, mutation on 101 would disrupt viral particle formation(Zhang et al., 2013a;Rouvinski et al., 2017). However, mD2VLP with a mutation of residue 101 can still form particles (Figure S13 in Supplementary Material), which further support our cryoEM model that the E:E protein interactions on VLP are loose. When the E:E protein interaction was looser, residue 101 at the groove within the dimeric molecule was more exposed (64%) than on the native virion (23.8%, PDB:3J27) or on the native virion during the 37°C “breathing” state (45.3%, PDB:3ZKO). We also generated a recombinant virus of DENV-2 whose EDIII was exchanged for a consensus EDIII(Chen et al., 2013) and the recombinant virus PL046cEDIII was recovered from the transfected cell culture supernatants for use in ELISA and FRμNT. Compared to the parental DENV-2 strain PL046, there was significant loss of binding of MAb DM25-3 and immune mouse sera to PL046cEDIII (Figure 6D), suggesting that DM25-3 could be an E-dimer inter-domain antibody whose binding footprint is sensitive to amino acid changes in the EDIII domain. By comparing FRμNT_50_ tests using murine antisera on both PL046 and PL046cEDIII viruses, we found that anti-mD2VLP sera showed a greater difference in neutralizing activity against PL046 and PL046cEDIII than did anti-imD2VLP sera, indicating a greater conformational dependence for anti-mD2VLP-triggered neutralization (Figure 6E).

Single B cells from the spleens of mD2VLP or imD2VLP-vaccinated mice were sorted into 96-well plates to better understand DENV-2 prM/E-specific B-cell repertoires (Figure 7A). By analyzing the amino acid sequences of the RT-PCR products of the variable regions of Ig heavy-chain (IgH) and light-chain (IgL) genes, the B-cell response from mD2VLP-immunized mice was more complex than the imD2VLP-immunized animals, particularly the gene loci of IgH (Figure 7B). The IgH and IgL genes from the DM25-3-producing hybridoma were also analyzed and found to contain IgHV1-22*01 and IgKV3-12*01 (Figure S14 in Supplementary Material), which were clonally expanded as shown in Figure 7B.

**Figure 7.**
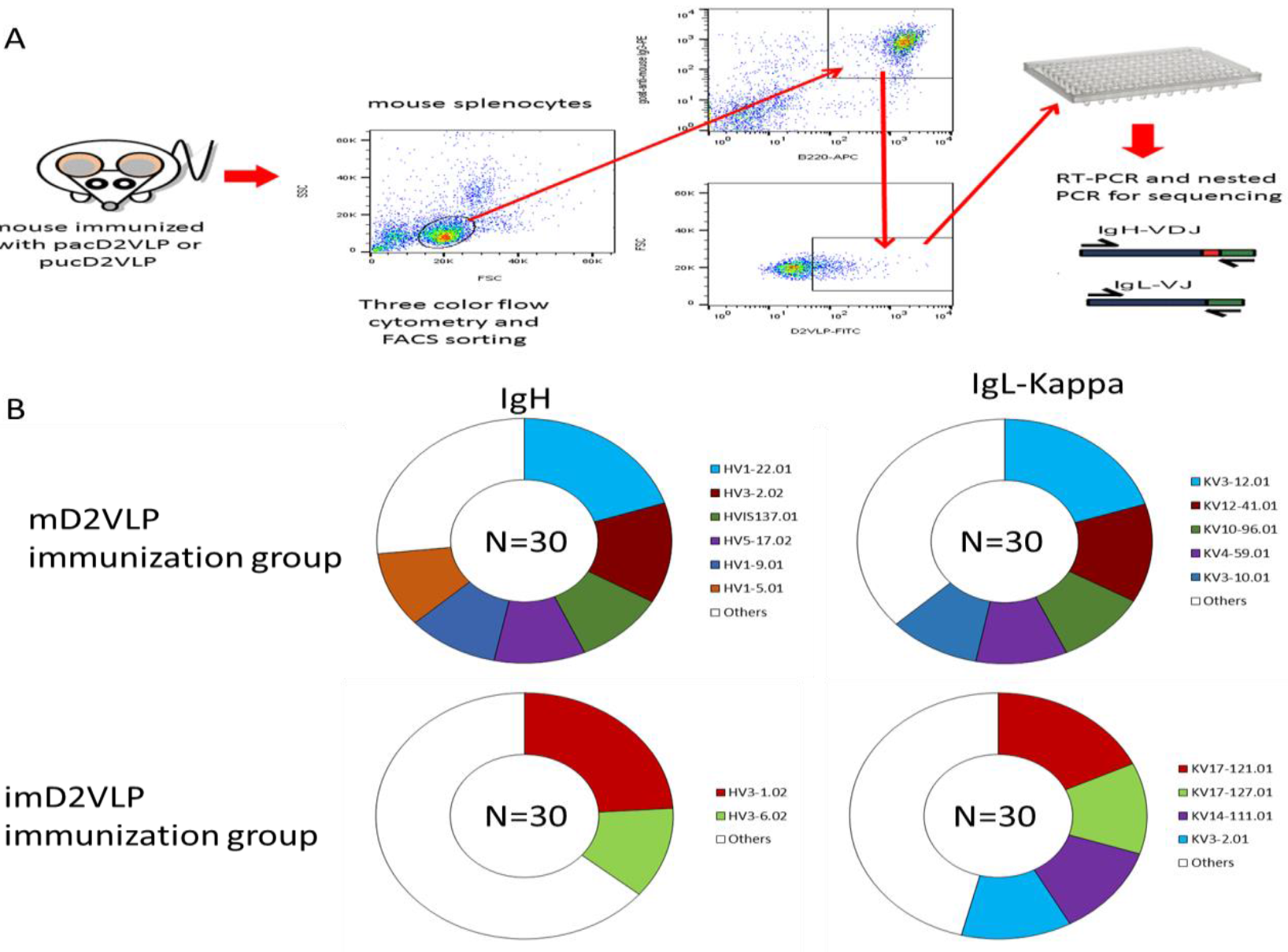
Overview of the experimental design and DENV-specific B-cell repertoires from the splenocytes obtained from mD2VLP- and imD2VLP-immunized mice. **(A)** Single DENV(+), B220(+) and IgG1(+)-specific B cell from the spleen of vaccinated mice was sorted into 96-well plates by FACS. The IgH and IgL gene transcripts of each single B cell were amplified by RT-PCR and nested PCR using gene-specific primers. **(B)** B-cell producing heavy chains and light chains encoded by the same IgH and IgL gene loci were grouped by the same color. The proportions of IgH- or IgL (κ-chain only)-genes are indicated by various colors corresponding to frequencies of the B-cell population of each vaccinated group. Frequencies greater than 10% are shown in various colors, otherwise they are grouped together and shown in white.

The major obstacle of dengue vaccine development is the lack of a suitable small animal model to evaluate vaccine efficacy. Therefore, we decided to test whether the monovalent mD2VLP immunogen could provide protection from a lethal dose challenge of heterologous dengue virus serotypes using suckling mice developed in a previous study(Chang et al., 2003;Hughes et al., 2012;Galula et al., 2014). The advantage of using sucking mice to test vaccine efficacy is that the protection from dengue virus challenge can only come from maternal antibodies generated by vaccinated female mice and passively transferred to the their babies. The immunization and challenge schedule is illustrated in Figure 8A. As shown in Figure 8B, suckling mice born from mothers vaccinated with mD2VLP were all protected from challenge with all four serotypes of dengue virus.

**Figure 8.**
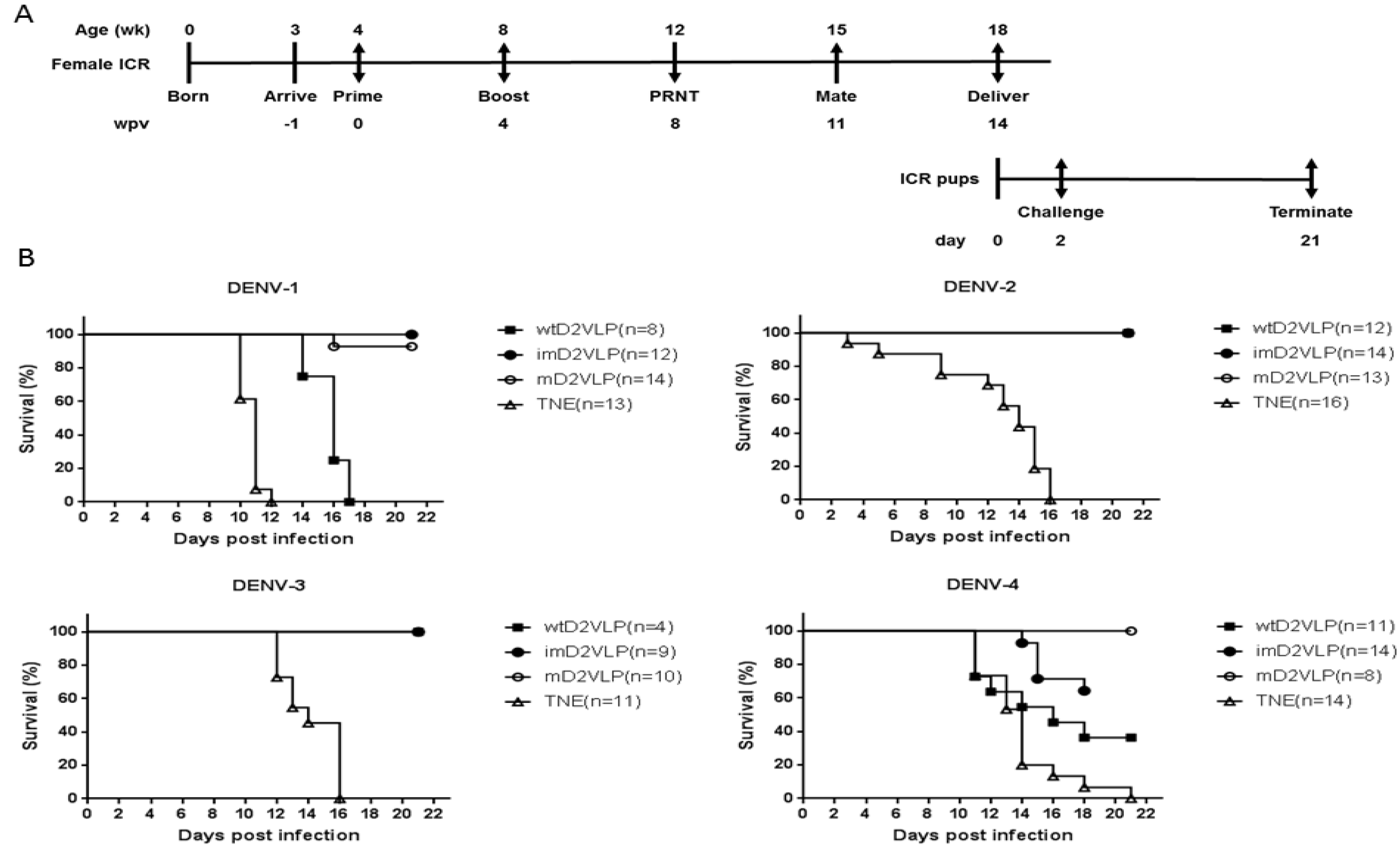
Schematic presentation of the schedule and survival curves for mouse immunization and challenge. **(A)** Groups of four 4-week-old, female, ICR mice were injected intramuscularly with imD2VLP and mD2VLP at week post vaccination (wpv) 0 and 4 at a dose of 4 pg/100 pL. Mice were bled from the retro-orbital sinus at week 4 following the second injection, and individual mouse serum collected from immunized females 1 week prior to mating was evaluated for the presence of the total IgG titer and the virus neutralization response by ELISA and focus-forming micro-neutralizing assay (FRNT). For the evaluation of passive protection by maternal antibody, ICR pups from the mating of non-immunized males with immunized females 11 weeks post initial vaccination were obtained for viral challenge. Pups from unvaccinated females were used as the challenge control. ICR pups from the designated groups were challenged individually through intracranial route at 2 days after birth with 10^4^ focus-forming unit (FFU) which were equivalent 141, 61, 11, 1000 times of 50% lethal doses (LD50) of DENV-1, to DENV-4, respectively. The percent survival of the mice was evaluated daily for up to 21 days. **(B)** Survival curve of pups delivered from the female mice receiving mD2VLP, imD2VLP monovalent vaccine or TNE control, then challenge with DENV-1 to 4 after birth. The in vivo protective efficacy of DENV-2 monovalent vaccine is maturity-dependent. N in parentheses indicated the numbers of pups of each group. Kaplan-Meier survival curves were analyzed by the log-rank test. ***, *P* <0.001.

## Discussions

How VLP immunogens mimic their viral counterparts structurally and how the neutralizing epitopes are preserved on the VLP surface are of significant interest in the development of good vaccine candidates. Recent studies suggest that potent human neutralizing antibodies with broad reactivity across dengue serocomplex can be generated from dengue patients, particularly after secondary infection(Tsai et al., 2013;Dejnirattisai et al., 2015;Rouvinski et al., 2015). Our unique findings here suggested that mD2VLP, with a scaffolding of multiple E-protein dimers similar to that of virion, is capable of inducing such CR antibodies with broad neutralizing activity. This monovalent mature-form dengue VLP with “epitope re-surfaced” has the potential to be the “epitope-focused” antigens when combined with other live attenuated dengue vaccine to induce higher and broader neutralizing antibodies(Rouvinski et al., 2017;Rey et al., 2018).

Very few studies explored the immunogenicity of prM-reduced antigens and its capability in inducing broad NtAbs(Keelapang et al., 2013;Suphatrakul et al., 2015;Metz et al., 2017;Rouvinski et al., 2017). Similar design of prM-reduced D2VLP was used as immunogens but was produced from mosquito cells and such VLP required adjuvants to boost immunity(Suphatrakul et al., 2015). Other attempts using E-dimers without the co-expression of prM cannot only display the quaternary structure epitopes but also reduce the exposure of FLE epitopes(Metz et al., 218;Metz et al., 2017;Rouvinski et al., 2017). However, the immunogenicity of such E-dimers with only two copies of E could be lower than VLP. The mD2VLP used in this study was produced from mammalian cells with the features similar to dengue virion, such as glycosylation pattern, particle distribution in gradient after high-speed centrifugation, epitopes preserved on the surface by the mapping of monoclonal antibodies, and mostly importantly, multiple copies of E structurally packed as dimers on the particle surface lattice (Fig S2 and S8 in Supplementary Material). Compared to imD2VLP (35%), only 7.6% of prM-recognizing antibodies were induced by mD2VLP through 2H2-blocking assay. Current dengue vaccine candidates in the market such as CYD-TDV or others in clinical trial are either prM-possessing forms or E-protein monomer which is lack of quaternary epitopes(Wichmann et al., 2017). Our study suggested that mD2VLP has the potential to be a safer immunogen as the second-generation dengue vaccine.

In addition, mD2VLP-vaccinated mice produce antisera predominantly composed of about 48% of DM25-3-like antibodies (Figure S15 in Supplementary Material). The packing of E dimers on VLPs does not just preserve quaternary structure epitopes on the surface lattice as suggested previously (Metz et al., 218;Crill et al., 2009), but also provides a unique surface-accessible structure with increased epitope accessibility. Usually these epitopes are cryptic in mature virions maintained at 28°C and in a neutral-pH environment(Lok, 2016). By superimposing the E-dimer-dependent quaternary epitopes(Lok et al., 2008;Fibriansah et al., 2015a;Rouvinski et al., 2015) onto the structure of mD2VLP, the binding footprints of these antibodies were highly overlapping and formed a “neutralization-sensitive hotspot” on mD2VLP (Figure S16 in Supplementary Material). Thus, the underlying mechanisms of why mD2VLPs are able to stimulate an elevated and broader immune response are based on the following: (1) epitope accessibility exposed at the grooves within E-protein dimers, which govern the generation of neutralizing antibodies; (2) removal of decoy epitopes presented on prM-containing structures found in imD2VLPs; and (3) inter-dimeric epitope accessibility due to the 5-fold and 3-fold openings of the E protein arrangement of T=1. Other mechanisms cannot be excluded such as reducing the production of FLE-like antibodies (ie. MAb E53), which have a preference of binding to spikes on noninfectious, immature flavivirions by engaging the highly conserved fusion loop that has limited solvent exposure of the epitope on mature virions(Cherrier et al., 2009).

Although the current obstacle in developing the second generation dengue vaccine is that dengue vaccine-induced neutralizing antibodies failed to correlate with or predict vaccine mediated protection(Moodie et al., 2018;Yang et al., 2018). However, studies of vesicular stomatitis virus demonstrated that *in vivo* protection depended on minimum serum antibody concentration, regardless of immunoglobulin subclass, avidity or in vitro neutralization activity(Bachmann et al., 1997). Therefore, it is crucial to understand the type of neutralizing antibodies induced by vaccination and establish the association between the level of such neutralizing antibodies and protection(Flipse and Smit, 2015;Henein et al., 2017;Katzelnick et al., 2017). Recent studies suggested that the most potent neutralizing antibodies came from those recognizing quaternary epitopes on the smooth surface of dengue virions(Fibriansah et al., 2015b;Rouvinski et al., 2015;Crowe, 2017), Regardless of the mechanisms involved in NtAb generation and protection from viral challenges, we have found that a mature form of monovalent VLP from dengue virus serotype 2 with “epitope re-surfaced” is efficient in inducing elevated and broadly NtAbs. When combined delivery with other live attenuated vaccine, such as CYD-TDV or dengue tetravalent vaccine, there is potential to provide not just T-cell immunity, but also higher broad NtAb response through “epitope-focusing” while exploring the next generation dengue vaccine in humans(Crill et al., 2012;Flipse and Smit, 2015;Rey et al., 2018).

Our unique findings in this study showed that a mature, monovalent DENV-2 VLP with grooves within the E protein dimeric molecules on the surface of particles induced highly protective, NtAbs against heterologous dengue viruses from all four serotypes. These characteristics strongly suggested this VLP with novel structure can combine with other live attenuated dengue vaccine as “immune-focused” antigens to induce higher and broader neutralizing antibodies. The strategy here may also provide a new direction for the development of other flavivirus vaccines, including Zika virus.

## Materials and Methods

### Ethics statement

This study was carried out in compliance with the guidelines for the care and use of laboratory animals of the National Laboratory Animal Center, Taiwan. The animal use protocol has been reviewed and approved by the Institutional Animal Care and Use Committee (IACUC) of National Chung Hsing University (Approval Number: 101-58). All efforts were made to minimize suffering of mice.

### Viruses, cells and antibodies

The strains of DENV serotypes 1-4 used were Hawaii (Genbank accession: KM204119.1), 16681 (Genbank accession: KU725663.1), H87 (Genbank accession: KU050695.1) and H241 (Genbank accession: AY947539.1), respectively. COS-1 (ATCC: CRL 1650), Vero (ATCC: CRL 1587) and C6/36 cells (ATCC: CRL 1660) were grown in Dulbecco’s modified Eagle’s medium (DMEM, Gibco, Life Technologies, Grand Island, NY) and were supplemented with 10% heat-inactivated fetal bovine serum (FBS, Hyclone, ThermoFisher, MA), 0.1mM nonessential amino acids (Gibco, Life Technologies, Grand Island, NY), 7.5% NaHCO3, 100U/ml penicillin, and 100ug/ml streptomycin; 5% FBS was used for Vero cells. Cells were maintained at 37°C with 5% CO_2_, except for C6/36 cells, which were maintained at 28°C without CO2. Parental DENV-2 strain PL046, generated from an infectious clone, and the domain III-swapped PL046 (PL046cEDIII), with domain III of DENV-2 replaced by consensus sequence as described previously (Liang et al., 2009;Chen et al., 2013), were provided by one of the coauthors, Dr. Y-L Lin.

Murine monoclonal antibody (MAb) 2H2, recognizing DENV prM protein, group cross reactive antibodies (4G2, 4A1B-9, 6B3B-3, 6B6C-1, 5H3) recognizing E-protein of all four major pathogenic flavivirus serocomplexes; complex cross-reactive antibodies (T5-1) recognizing all four DENV serocomplex viruses; serotype-specific MAb 3H5-1, recognizing DENV-2 only; rabbit sera and mouse hyper-immune ascitic fluid (MHIAF) for DENV-1 to 4 were provided by one of the authors, G-J Chang (DVBD, CDC, Fort Collins, CO). DENV-2 E protein-specific MAb 32-6 was provided by Dr. H-C Wu (Academia Sinica, TW), while DENV-2 prM-specific MAb 155-49 was obtained from H-Y Lei (National Cheng Kung University, Taiwan). MAb 2H2 was also labeled with biotin using EZ-Link™ Sulfo-NHS-Biotin kit (Thermo Fisher Scientific Inc., Rockford, IL) according to the manufacturer’s instructions. In order to increase the recognition of M protein, anti-M antibody was produced by cloning the complete M protein sequence into pET21a (Novagen, Germany) and then purifying the expressed protein under denaturing conditions as described in the molecular cloning handbook. Ten micrograms (10μg) of purified M protein with Freud’s complete adjuvant was used to immunize the mice by the intraperitoneal route five times at 2-week intervals. Anti-mouse CD45R/B220 antibody conjugated with APC (allophycocyanin) and goat anti-mouse IgG conjugated with PE (phycoerythrin) were purchased from BioLegend (San Diego, CA).

Previously constructed and characterized recombinant plasmid pVD2, expressing the prM, 80% and 20% COOH terminus of the envelope proteins of DENV-2 (Asian 1 genotype, strain 16681) and Japanese encephalitis virus (strain SA14-14-2), respectively (Galula et al., 2014), was used in this study. The furin cleavage site of prM was mutated in pVD2 to generate a prM-uncleaved plasmid or as indicated in Figure 1A by using site-directed mutagenesis following the manufacturer’s protocol (Stratagene, La Jolla, CA). The primers used for cloning and site-directed mutagenesis are provided in Table S1. Nucleotide sequencing confirmed that all the plasmids contained no other mutations other than those indicated.

### Sequence analysis of flavivirus prM sites

Flavivirus prM protein amino acid sequence alignments were performed using ClustalX 2.1 software with the representative strains and the GenBank accession numbers as follows: dengue virus serotype 2 (NP_056776), dengue virus serotype 1 (AIU47321), dengue virus serotype 3 (YP_001621843), dengue virus serotype 4 (NP_073286), Japanese encephalitis virus (NP_775664), St. Louis encephalitis virus (AIW82235), West Nile virus (AI010814), tick-borne encephalitis virus (NP_775501), yellow fever virus (NP_041726), cell-fusion agent virus (NP_041725), Zika virus (BAP47441.1). Sequences of prM junction region were scored for their predicted cleavability by furin using the PiTou 2.0 software package(Tian et al., 2012). A negative score indicates a sequence predicted not to be cleaved by furin, whereas a positive score denotes prediction of furin cleavability.

### Antigen production and purification

To produce virus-like particle (VLP) antigens, COS-1 cells at a density of 1.5×10^7^ cells/mL were electroporated with 30 μg of each pVD2 plasmid following the previously described protocol (Chang et al., 2000). After electroporation, cells were seeded into 75-cm^2^ culture flasks (Corning Inc., Corning, NY, USA) containing 15 mL growth medium and allowed to recover overnight at 37°C. The growth medium was replaced the next day with a maintenance medium containing serum-free medium (SFM4MegaVirTM, SH30587.01, Hyclone, ThermoFisher) supplemented with non-essential amino acid (NEAA), GlutaMAX, sodium pyruvate and cholesterol (Gibco, Life Technologies, Grand Island, NY), and cells were continuously incubated at 28°C with 5% CO_2_ for VLP secretion. Tissue-culture media were harvested 3 days after transfection and clarified by centrifugation at 8,000×g for 30 minutes (mins) at 4°C in AF-5004CA rotor (Kubota, Tokyo, Japan) using a Kubota 3740 centrifuge. The harvested media were first concentrated 20-fold using 100K Amicon Ultra centrifugal filters (Merck Millpore, CA) before loading onto a 20% sucrose cushion and concentrated by ultracentrifugation at 28,000 rpm for 16 hours at 4°C in a Beckman SW28 rotor. Purified VLPs were resuspended in 250μl TNE buffer (50mM Tris-HCl, 100mM NaCl, 0.1 mM EDTA, pH 7.4) for every liter of harvested medium at 4°C overnight. The VLPs were further purified by rate zonal centrifugation in a 5 to 25% sucrose gradient at 25,000 rpm at 4°C for 3 hours. All gradients were made with TNE buffer and were centrifuged in a Beckman SW41 rotor. Fractions of 0.5 ml were collected by upward displacement and assayed by antigen-capture ELISA. For experiments that required highly purified VLPs, proteins with the peak OD values from antigen-capture ELISA were pelleted at 40,000 rpm at 4°C for 4 hours using a Beckman SW41 rotor, and re-suspended in 250μl TNE buffer. The protein concentration was measured by the Bradford assay (BioRad, Hercules, CA) following the commercial protocol and using bovine serum albumin (BSA, New England Biolabs, MA) as a standard. Purified VLPs were also labeled with fluorescein for antigen-specific B-cell sorting following the manufacture’s protocol (Zhang et al., 2010).

### ELISA

The ratio of prM to E protein was determined using DENV-2 immune rabbit serum to capture VLPs. The DENV-2 E and prM proteins were measured by ELISA using MAb 3H5 (specific for DENV-2 domain III) and MAb 155-49 (specific for DENV prM). The ratio was calculated as absorbance for prM/absorbance for E protein. The uncleaved prM VLP (imD2VLP), which contained amino acid mutations at P1 and P2 sites from amino acid residue R/K to T/S (Li et al., 2008), respectively, was used as a standard to calculate the percentage of prM cleavage. Percent cleavage of prM was then calculated with reference to imD2VLP, which was assumed to be 100% uncleaved, as previously described (Dejnirattisai et al., 2015).

Antigen-capture ELISA was performed to quantify the amount of different VLP antigens. Briefly, flat-bottom 96-well MaxiSorp™ NUNC-Immuno plates (NUNC™, Roskilde, Denmark) were coated with 50 μL of rabbit anti-DENV-2 VLP serum at 1:500 in bicarbonate buffer (0.015 M Na_2_CO_3_, 0.035 NaHCO_3_, pH 9.6), incubated overnight at 4°C, and blocked with 200 μL of 1% BSA in PBS (1% PBSB) for 1 hr at 37°C. Clarified antigens were titrated two-fold in PBSB, and 50 μL of each dilution was added to wells in duplicate, incubated for 2 hrs at 37°C, and washed five times with 200 μL of 1x PBS with 0.1% Tween-20 (0.1% PBST). Normal COS-1 cell tissue culture fluid and cell pellets were used as control antigens. Captured antigens were detected by adding 50 μL of anti-DENV-2 MHIAF at 1:2,000 in blocking buffer, incubated for 1 hr at 37°C, and washed for five times. Fifty microliters of HRP-conjugated goat anti-mouse IgG (Jackson ImmunoResearch, Westgrove, PA, USA) at 1:5,000 in blocking buffer was added to wells and incubated for 1 hr at 37°C to detect MHIAF. Subsequently, plates were washed ten times. Bound conjugate was detected with 3,3’,5,5’-tetramethylbenzidine substrate (Enhanced K-Blue^®^ TMB, NEOGEN^®^ Corp., Lexington, KY, USA), after incubation at room temperature for 10 mins, and addition of 2N H_2_SO_4_ to stop the reaction. Reactions were measured at A_450_ using a Sunrise™ TECAN microplate reader (Tecan, Grödig, Austria). The capability of antigen-capture ELISA to detect imD2VLP or mD2VLP was measured against purified D2VLP at known total protein concentration from individual preparations. Data are expressed as P/N ratio by dividing the OD450 value from each dilution of mouse sera or MAb by the OD450 value from the control COS-1 culture supernatant.

An IgG ELISA was used to assay the presence of antigen-specific IgG in the postvaccination mouse sera using the same antigen-capture ELISA protocol described above with minor modifications. Equal amounts of purified antigens were added into each well of an Ag-capture ELISA plate. The concentration of purified antigens were determined from the standard curves generated using a sigmoidal dose-response analysis using GraphPad Prism (version 6.0, GraphPad Software, Inc., La Jolla, CA, USA). Individual mouse sera, collected from mice which had the same immunization schedule, initially diluted at 1:1,000, were titrated two-fold and added into wells in duplicate, and were incubated for 1 hr at 37°C. Pre-vaccination mouse sera were used as negative controls. Incubations with conjugate and substrate were carried out according to the standard Ag-capture ELISA as outlined. The OD_450_ values, modeled as nonlinear functions of the log10 serum dilutions using a sigmoidal dose-response (variable slope) equation and endpoint antibody titers from two independent experiments, were determined as the dilutions where the OD value was twice the average OD of a negative control.

Epitope-blocking ELISAs were performed to determine the vaccinated mouse response to the prM protein. The setup was similar to the IgG ELISA wherein plates were coated with rabbit anti-DENV-2 VLP serum, and blocked with 1% BSA in PBS. After washing, pooled mouse sera were diluted two-fold in blocking buffer starting from 1:1,000, and were then incubated with imD2VLP antigen (pre-titrated to OD_450_=1.0) for 1 hr at 37°C. After serum incubation and washing, MAb 2H2 conjugated with biotin by EZ-Link Sulfo-NHS-Biotin (ThermoFisher, CA) at 1:4000 dilution was added to each well and incubated for 1 hr at 37°C to compete with the already-bound antibody from the immune mouse sera specific for the imD2VLP antigen. Bound MAb 2H2 conjugate was detected with 1:1000 HRP-conjugated streptavidin (016-030-084, Jackson ImmunoResearch), and was incubated for 1 hr at 37°C. After washing with PBS for ten times, TMB substrate was added into the wells and the plates were incubated for 10 mins, and the reaction was stopped with 2 N H_2_SO_4_. Reactions were measured at A_450_. Percent blocking was determined by comparing replicate wells with Biotin-conjugated MAb competing against pre-adsorbed naïve mouse serum using the formula: % Blocking = [OD450 of imD2VLP-OD_450_ of imD2VLP blocked by MAb 2H2)/OD_450_ of imD2VLP]×100.

Binding-ELISAs were used to assess the binding activity of MAbs or mouse immune sera to D2VLP or mutant antigens using a similar antigen-capture ELISA set-up, except that two-fold dilutions of the specific MAb or immune mouse sera replaced the anti-DENV-2 MHIAF. Equal amounts of D2VLP antigens were added into wells, and were standardized using purified D2VLPs. The antibody endpoint reactivity was determined in a similar manner to the determination of antigen endpoint secretion titers.

### SDS-PAGE and Western blotting

Equal amounts of purified VLPs and sample buffer were mixed and analyzed by 12% Tricine-sodium dodecyl sulfate-polyacrylamide gel electrophoresis (Tricine-SDS-PAGE)(Schägger, 2006). For immunodetection, proteins were blotted from gels onto nitrocellulose membranes (iBlot^®^2NC mini stacks, ThermoFisher Scientific) with iBlot^®^ 2 Gel Transfer Device (ThermoFisher Scientific). The membranes were incubated for 1 hour at room temperature in phosphate-buffered saline (pH 7.4) containing 5% skim milk (BD biosciences, CA) to block nonspecific antibody binding. After 1hr incubation, membranes were individually stained to detect E, prM and M proteins by using anti-DENV2 MHIAF at 1:2000, anti-DENV prM MAb 155-49 at 0.5μg/ml and mouse anti-M sera at 1:25, respectively, at 4°C overnight. Membranes were washed three times for 15 min each. DENV-specific bound immunoglobulin was recognized with HRP-labeled goat anti-mouse IgG (Jackson ImmunoReasearch, PA), and was visualized with ECL (enhanced chemiluminescent substrate, GE Healthcare, UK) according to the manufacturer’s protocol. The ECL signals were detected by ImageQuant™ LAS 4000 mini (GE Healthcare). However, M protein was visualized using TMB membrane peroxidase substrate (KPL, MD) to avoid high background from ECL.

### Mouse experiment

Groups of four 4-week-old female BALB/c mice were injected intramuscularly with imD2VLP and mD2VLP at weeks 0 and 4 at a dose of 4 μg/100 μL PBS divided between the right and left quadriceps muscle. Mice were bled from the retro-orbital sinus at week 4 following the second injection, and individual serum specimens were evaluated for DENV-2 specific antibodies by ELISA and focus-reduction micro-neutralization test (FRμNT), as described in the following section.

For the evaluation of passive protection by maternal antibody, ICR pups from the mating of non-immunized males with immunized females 11 weeks post initial vaccination were obtained for viral challenge. Immune sera were collected from immunized females 1 week prior to mating to confirm the presence of the total IgG titer as well as the virus neutralization antibody titer. Pups from unvaccinated females were used as the challenge control. ICR pups from the designated groups were challenged individually through the intracranial route at 2 days after birth with 10^4^ focus-forming units (FFU) which were equivalent to 141, 61, 11, 1000-fold of 50% lethal doses (LD50) of DENV-1 (strain Hawaii), DENV-2 (strain 16681), DENV-3 (strain H87) and DENV-4 (strain BC71/94, kindly provided by one of the co-author, Dr. Chang from US-CDC), respectively. Mouse survival was evaluated daily for up to 21 days.

### Virus neutralization

The neutralizing ability of the immune mouse sera for all serotypes of DENV was measured by focus-reduction micro neutralization test (FRμNT), as previously described(Galula et al., 2014). Briefly, 2.475 × 10^4^ Vero cells/well were seeded onto flat-bottom 96-well Costar^®^ cell culture plates (Corning Inc., Corning, NY, USA) and incubated for 16 hrs overnight at 37°C with 5% CO_2_. Pooled sera were initially diluted at 1:10, heat-inactivated for 30 min at 56°C, titrated two-fold to a 40 μL volume, and 320 pfu/40 μL of DENV-1 to 4 was added to each dilution. The mixtures were then incubated for 1 hr at 37°C. After incubation, 25 μL of the immune complexes were added in duplicates onto plates containing Vero cell monolayers. Plates were incubated for 1 hr at 37°C with 5% CO_2_ and rocked every 10 mins to allow infection. Overlay medium containing 1% methylcellulose (Sigma-Aldrich Inc., St. Louis, MO, USA) in DMEM with 2% FBS was added, and plates were incubated at 37°C with 5% CO_2_. Forty-eight hours later, plates were washed, fixed with 75% acetone in PBS and air-dried. Immunostaining was performed by adding serotype-specific MHIAF at 1:600 in 5% milk and 0.1% PBST and incubated for 60 min at 37°C. Plates were washed and goat anti-mouse IgG-HRP at 1:100 in 5% milk and 0.1% PBST was added; plates were incubated for 45 min at 37°C. Infection foci were visualized using a peroxidase substrate kit, Vector^®^ VIP SK-4600 (Vector Laboratories, Inc., Burlingame, CA, USA), following the manufacturer’s instructions. FRμNT titers were calculated for each virus relative to a virus only control back-titration. Titers were determined as a 50% reduction of infection foci (FRμNT50) and were modeled using a sigmoidal dose-response (variable slope) formula. All values were determined from the average of two independent experiments. Target virus strains were: DENV-1, Hawaii; DENV-2, 16681; DENV-3, H87; and DENV-4, H241. For E-dimer, inter-domain-specific neutralizing antibodies, the recombinant parental DENV-2 strain PL046 and EDIII-swapped PL046cEDIII were used as target viruses. In the calculation of geometric mean titers (GMT) for graphic display and statistical analysis, a FRμNT50 titer of <10 was represented with the value of 1 and 5, respectively.

### Generating hybridomas and MAb screening

Hybridomas secreting anti-DENV antibodies were generated from the mD2VLP immunized mice according to a standard procedure(Kohler and Milstein, 1975), with slight modifications(Chen et al., 2007). First, the mD2VLP-immunized mouse was boosted with another 4μg of mD2VLP 24 weeks prior to terminal bleeding. At day 4 after the third immunization, splenocytes were harvested from the immunized mouse and fused with NSI/1-Ag4-1 myeloma cells using an antibody delivery kit following manufacturer’s recommendations (GenomONETM-CF HVJ Envelope Cell Fusion Kit, Gosmo Bio Co, ISK10 MA17). Fused cell pellets were suspended in DMEM supplemented with 15% FBS, hypoxanthine-aminopterin-thymidine medium, and hybridoma cloning factor (ICN, Aurora, OH). Hybridoma colonies were screened for secretion of MAbs by ELISA following the procedures as described above. However, the cell culture supernatant of C6/36 cells infected with DENV-2 virus at a multiplicity of infection (moi) equal to 1.0 was used as the antigens and the culture supernatant from each of the hybridoma colonies was used as the detecting antibody. Selected positive clones were subcloned by limiting dilution. Ascitic fluids were produced in pristane-primed BALB/c mice. Hybridoma cell lines were grown in DMEM with 10% heat inactivated FBS. MAbs were affinity purified with protein G Sepharose 4B gel, and the amount of each purified MAb was quantified by comparison with a known amount of IgG used in a standard ELISA.

### Profiling of prM/E-specific Ig genes from a single B-cell

Ig genes from a single B cell were isolated following the protocols established by previous reports with some modifications(Tiller et al., 2009). Briefly, single cell suspensions were collected from the spleens of BALB/c mice immunized twice at 4-week intervals with imD2VLP and mD2VLP. The D2VLP(+), B220 (+) and IgG1(+) splenic B cells were isolated by flow cytometry (FACSAria II), and single B cells were sorted into 96-well PCR plates containing 4ul/well of ice-cold RNase-free water supplemented with 10mM DTT and 3U RNase inhibitor (Promega). RT-PCR reactions were performed using the Qiagen OneStep RT-PCR kit (Qiagen) at 50°C for 30 mins and at 95°C for 15 min followed by 40 cycles of 95°C for 1 min, 55°C for 1 min and 72°C 1 min, and a final incubation at 72°C for 5 min, with the primers as previously described. The nested PCR was performed using 2 ul of unpurified, first-round PCR product at 95°C for 1 min followed by 40 cycles of 95°C 30 sec, 57°C (IgH) or 45°C (Igk) or 55°C (Igλ) for 30sec, 72°C for 45sec, and incubated at 72°C for 10 min as previously described. Aliquots of nested PCR products were sequenced and analyzed using IMGT/V-Quest (http://www.imgt.org) to identify the highest homology gene loci of germ-line V, D and J genes. Ig genes were then translated and aligned by CLUSTALW to define the clonally amplified Ig genes.

### Immunogold labeling and transmission electron microscopy (TEM)

To detect mature DENV-2 VLPs (mD2VLPs) as spherical particles, 3μL of freshly prepared samples were adsorbed onto a glow-discharged nickel grid (EMS CF-200-Ni) for 20 mins, The grid containing the samples was incubated in 1% BSA/PBS blocking buffer for 1 hour at room temperature. As soon as the excess liquid was removed with a filter paper, DENV-2-specific MAb DB32-6 was used as the primary antibody (dilution 1:500), and was added to the samples for 2 hrs at room temperature. Then the grid was washed with blocking buffer and incubated with 6-nm nanogold-conjugated Donkey anti-mouse IgG (Abcam, MA) (dilution 1:30) at room temperature for 1 hr. The excess liquid was removed with filter paper and fixed with 1% glutaraldehyde in PBS buffer (GA/PBS) for 10 mins. After fixing, excess liquid was removed using a filter paper. Finally, the sample was stained with 2% uranyl acetate (UA) for 1 min and air-dried. The immunogold labeling process was performed in a high humidity chamber. The immunogold labeled mD2VLPs were inspected by TEM. The images were taken by a JEM1400 electron transmission microscope at a magnification of 100,000x using a 4k × 4k Gatan 895 CCD camera. The diameters of particles were measured by ImageJ software.

### Cryo-EM and 3D reconstruction

A freshly prepared dengue virus sample (3 μl) was placed onto a glow-discharged Quatifoil 2/2 grid (Quatifoil GmbH, Germany), blotted with filter paper, and plunged into liquid nitrogen-cooled liquid ethane using Gatan CP3. Cryo-EM images were recorded with a JEM2100F using an accelerating voltage of 200kV and a magnification of 15,000x using a direct electron detector (DE-12 Camera System - Direct Electron, LP) with a 6 μm pixel size (corresponding to ~4 Å at the specimen level). The measured defocus values of these images range from −2μm to −4.5μm. The imaging electron dosage was ~ 10 e^−^/Å^2^.

Variations in size and shape were observed in the EM images (Figure S3A, B in Supplementary Material). The irregularly-shaped particles were eliminated through visual inspection, while the spherical particles were subjected to further image analyses. The particle size analyses showed that the major peaks were located at ~26nm, ~31nm and ~36nm diameter size classes which were therefore denoted by small (‘S’), medium (‘M’) and large (‘L’). The heterogeneity analyses by EMAN2(Tang et al., 2007) and XMIPP(Sorzano et al., 2004;de la Rosa-Trevín et al., 2013) showed that all of the classes had distinct two layers and that the particles classes with diameters of 31nm had more prominent features than others (Figure S3B, C in Supplementary Material). Taken together with the immunogold labeling results, we selected the particles with size of ~31nm or further 3D reconstruction process. The structure reconstruction processes were performed by EMAN2. There were 4,217 spherical particles with the size of ~31nm which were included in the reconstruction process and the icosahedral symmetry was imposed during the whole process. The reconstruction process ended when there was no improvement achieved. The resolution of the final reconstructions was 13.1Å from a Fourier shell correlation curve using the gold standard resolution estimate in EMAN2 (Figure S5 in Supplementary Material). Solvent accessible surface area (SASA) of individual amino acid molecules on mD2VLP was calculated by POPS program(Cavallo et al., 2003;Fornili et al., 2012).

### Fitting

The atomic structure of mD2VLP was modeled based on the cryo-EM structure of the mature dengue virus at 3.5Å resolution (PDB ID: 3J27) using MODELLER 9v15(Eswar et al., 2007). The envelop proteins were fitted rigidly using the Fit-in-Map tool in *UCSF Chimera*, when the CCC (Cross-Correlation Coefficient) score was maximized, the complete copies were generated to follow the T=1 arrangement with *Multiscale Models* using Icosahedral symmetry, xyz 2-fold axes (VIPER).

### Statistical analysis

All data are represented as means ± standard error, and analyzed using GraphPad Prism version 6.0. Unpaired t test was used to analyze data sets between two groups. Mann-Whitney U test to account for non-normality of some transformed data was also applied. P values <0.05 were considered significant.

## Data and materials availability

The cryo-EM density map of mD2VLP has been deposited to Electron Microscopy Data Bank under accession number EMDB6926. All other relevant data are available from the authors on request.

## Acknowledgments

We thank Dr. Felix Rey for commenting on the structure of dengue VLP and Ann Hunt for English editing.

## Author contributions

WFS performed the characterization of D2VLP including Western blotting, ELISA, site-directed mutagenesis and mouse immunization. JHL conducted the Pitou and solvent accessibility analysis. MYL, DYC, HCW and JUG produced and characterized the NS1-splenocytes fused B cell lines, monoclonal antibody and single B-cell repertoire characterization. WFS and YCW conducted VLP purification, mouse immunization experiments and neutralization assays. JUG and CHH prepared the anti-M protein mouse polyclonal serum. JJL and YLL prepared the parental and domain III-replaced recombinant DENV-2. SRC and SRW prepared the cryo-EM samples, processed the cryo-EM images, fitted the protein structures, and solved the cryo-EM structure. MTW, GJC and WFS mapped the 2H2 binding epitopes. SRW, GJC and DYC supervised the studies and contributed to the writing of the paper.

## Competing interests

All the authors declare that they have no competing interests.

## Funding

This study was supported by Ministry of Science and Technology Taiwan (MOST 104-2320-B-006-027 and MOST 105-2320-B-006-017-MY3).

## Supplementary Materials

**Table S1.**
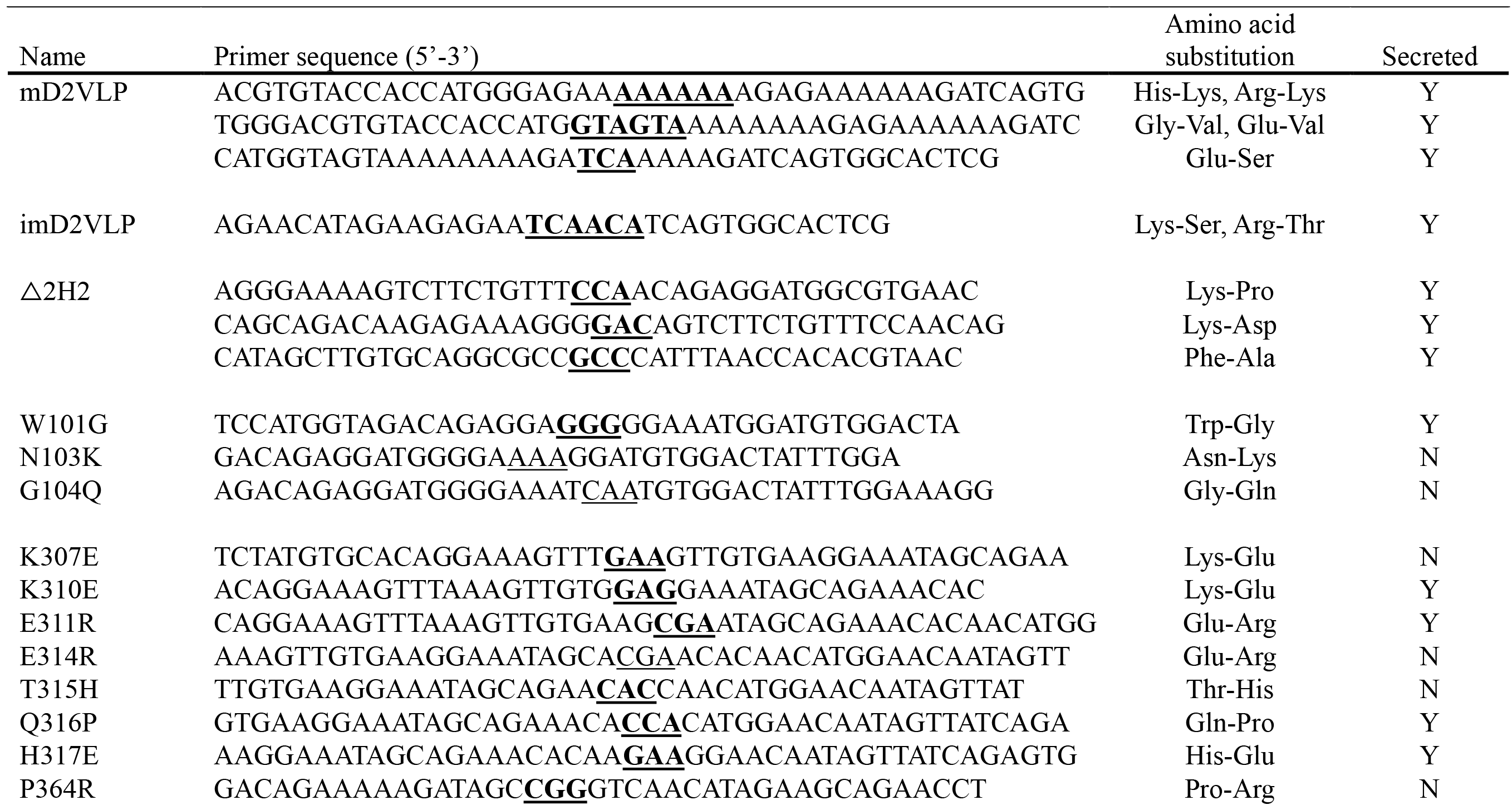
Nucleotide sequences of primers for site-directed mutagenesis used in this study

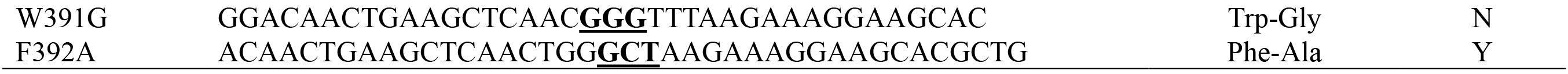

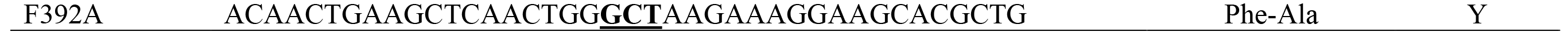

**Figure S1.**
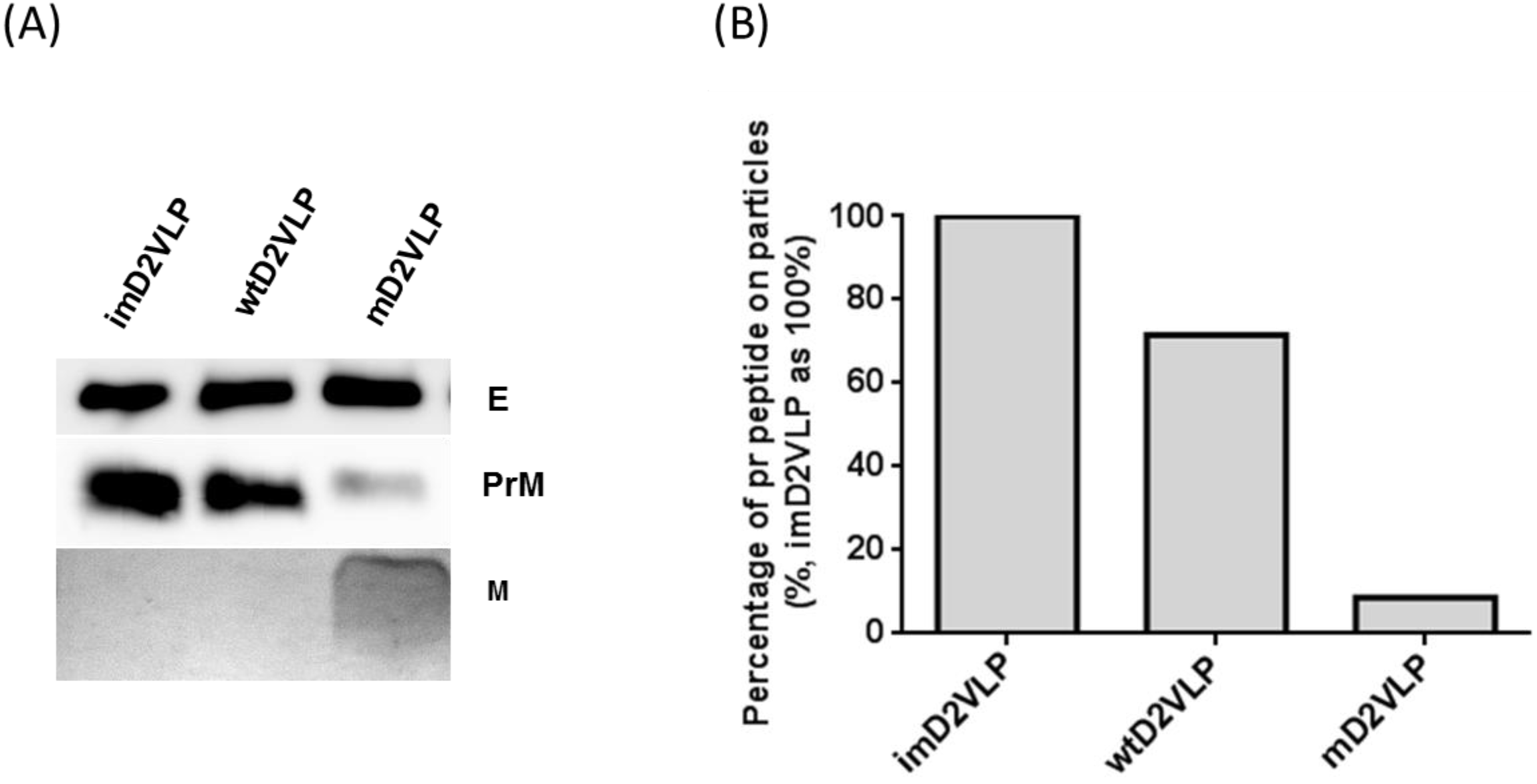
Comparison of prM cleavage among different DENV-2 virus-like particles (D2VLP). **(A)** Culture supernatants of mD2VLP and imD2VLPs were collected and purified after electroporation with the respective plasmids. Five micrograms of proteins were loaded onto a 12% non-reducing Tricine-SDS-PAGE. E, prM and M proteins were assayed by Western blot using mouse hyper-immune ascitic fluids (MHIAF, 1:2000), MAb 2H2 (0.5 μg/mL) and anti-M protein mouse sera (1:25), respectively. E and prM proteins were visualized with enhanced chemiluminesence (ECL); however, M protein was visualized by TMB substrate to avoid high background. **(B)** Percentages of pr-peptide remaining on particles were measured by calculating the relative densitometric quantity of prM band by Bio-1D software.

**Figure S2.**
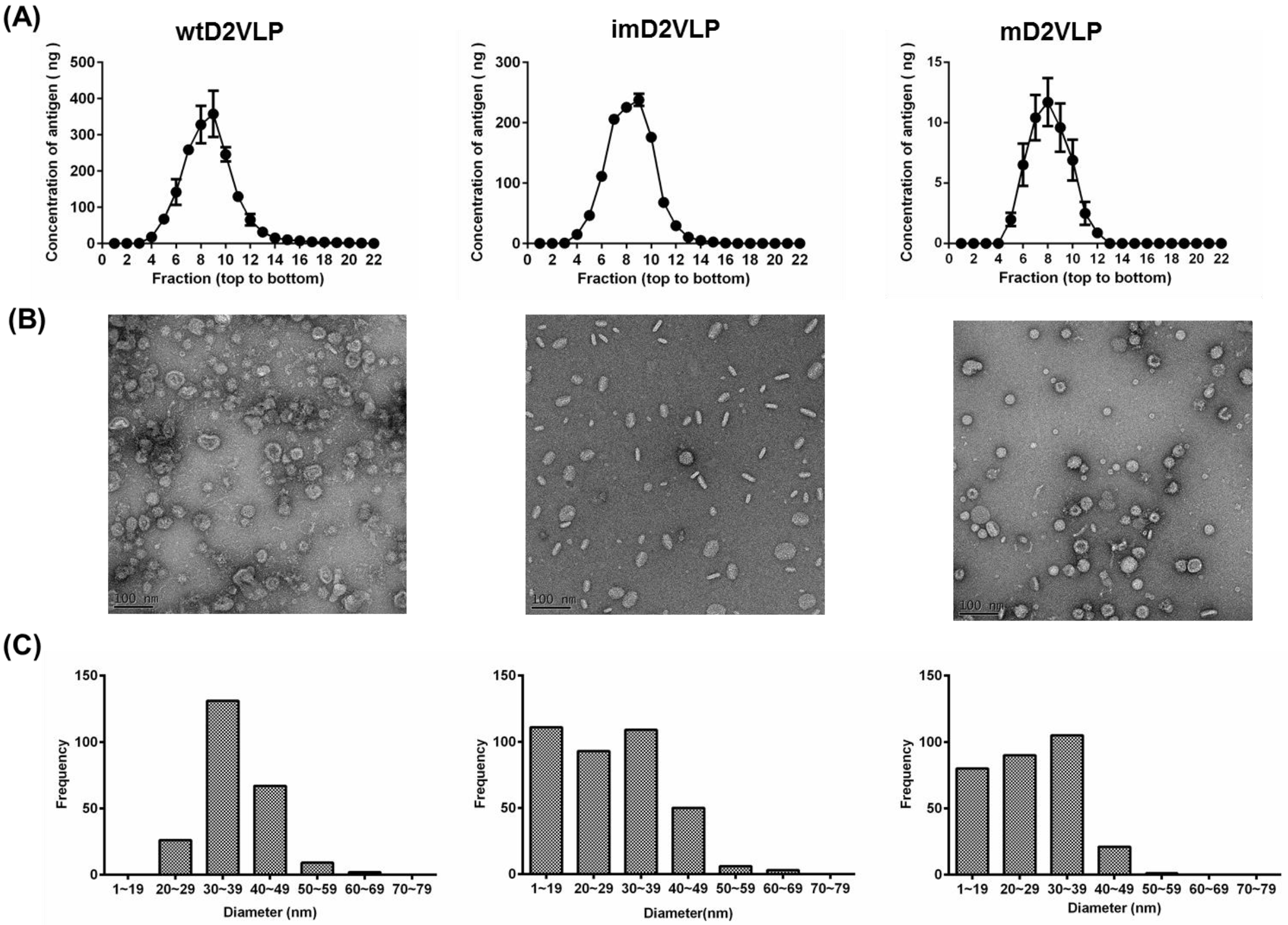
Comparison of physical properties of D2VLP among different DENV-2 virus-like particles (D2VLP). (A) Equilibrium banding profiles of wtD2VLP (left), imD2VLP (center) and mD2VLP (right) after rate-zonal centrifugation on a 5-25% linear sucrose density gradient at 25,000 rpm for 3 hours. Equivalent amounts (50μl) of D2VLP from each fraction were subjected to antigen-capture ELISA. (B) Electron micrographs at 50,000-fold magnification of wtD2VLP (left), imD2VLP (center) and mD2VLP (right) stained with uranyl acetate from the peak fraction in (A). (C) The diameter of 200 randomly selected particles of wtD2VLP (left), imD2VLP (center) and mD2VLP (right) from representative electron micrographs as in (B) were determined by software Gatan Digital Micrograph. (D) Endoglycosidase treatment ofpurified wtD2VLP, imD2VLP and mD2VLP. Samples were treated with PNGase F or Endo H and compared with untreated controls by 10% non-denaturing SDS-PAGE and immunoblotting with DENV-2 mouse hyper-immune ascitic fluid (MHIAF).

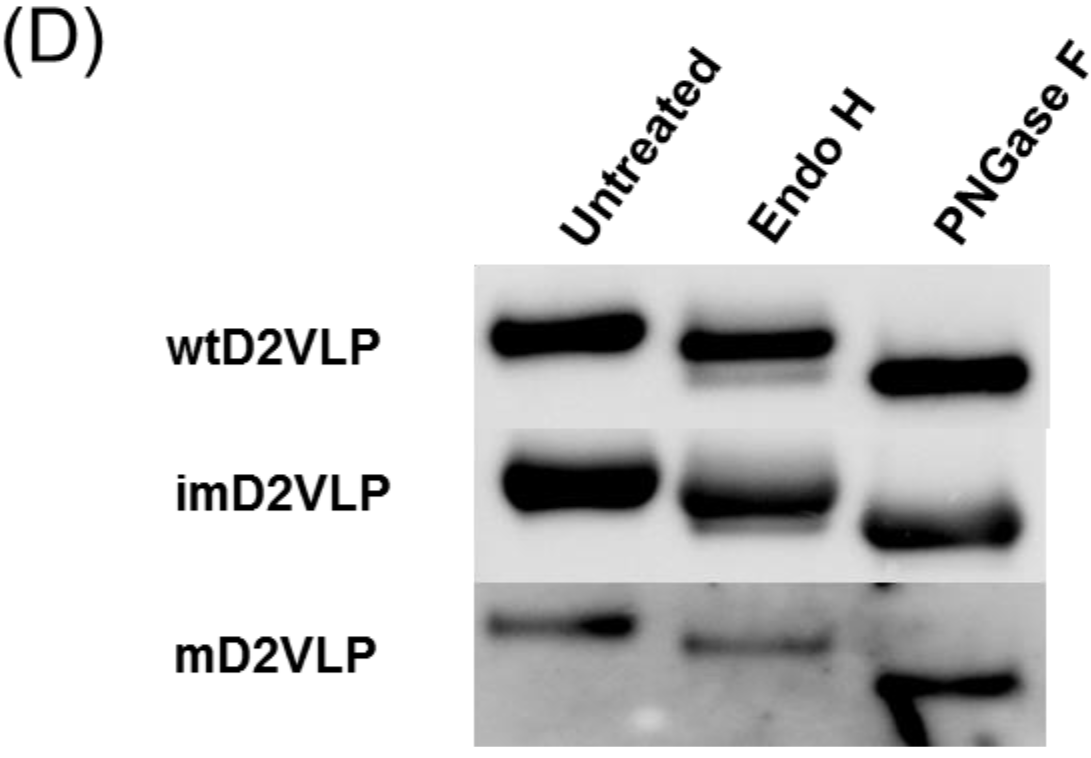

**Figure S3.**
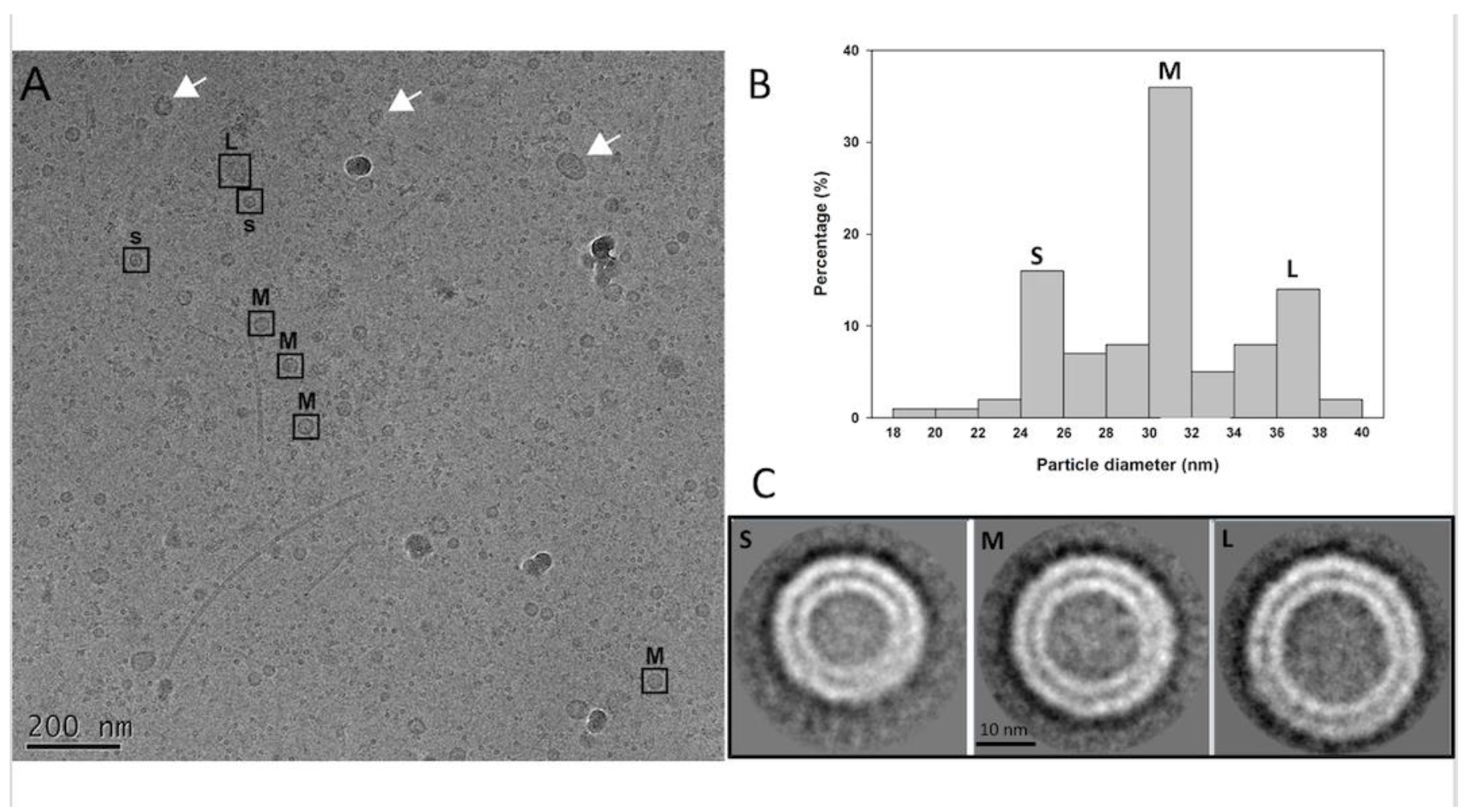
The cryo-EM images and 2D analysis of the particles. (A) Cryo-EM images of purified mD2VLP showed spherical particles (boxed) among irregular or incomplete structures (arrows). Irregularly-shaped particles were eliminated through visual inspection, while the spherical particles were subjected to further image analyses. (B) The particle size analyses showed the size variation in the sample and that the major peaks were located at ~26nm, ~31nm and ~36nm diameter size classes which were therefore denoted by small (‘S’), medium (‘M’) and large (‘L’). Each of the ‘S’, ‘M’ and ‘L’ particles identified was labeled. (C) The 2D image analyses showed that the particles had two distinct layers and that the particles classes with diameters of ~31nm had more solid features than others.

**Figure S4.**
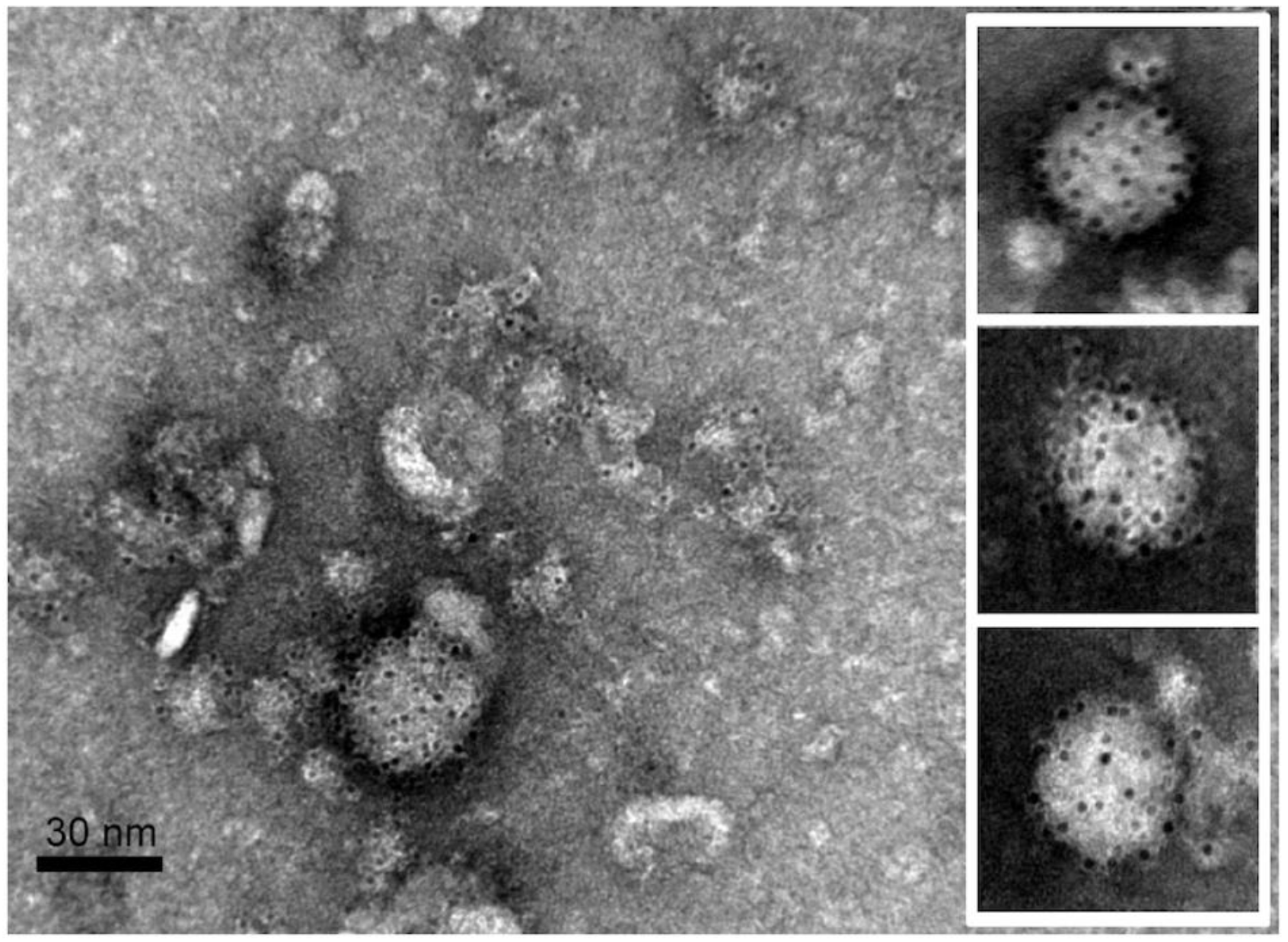
The EM images of immunogold-labeled mD2VLPs. The E proteins were labeled with domain III-specific monoclonal antibody (MAb 32-6), and 1113 conjugated with 6-nm gold particles

**Figure S5.**
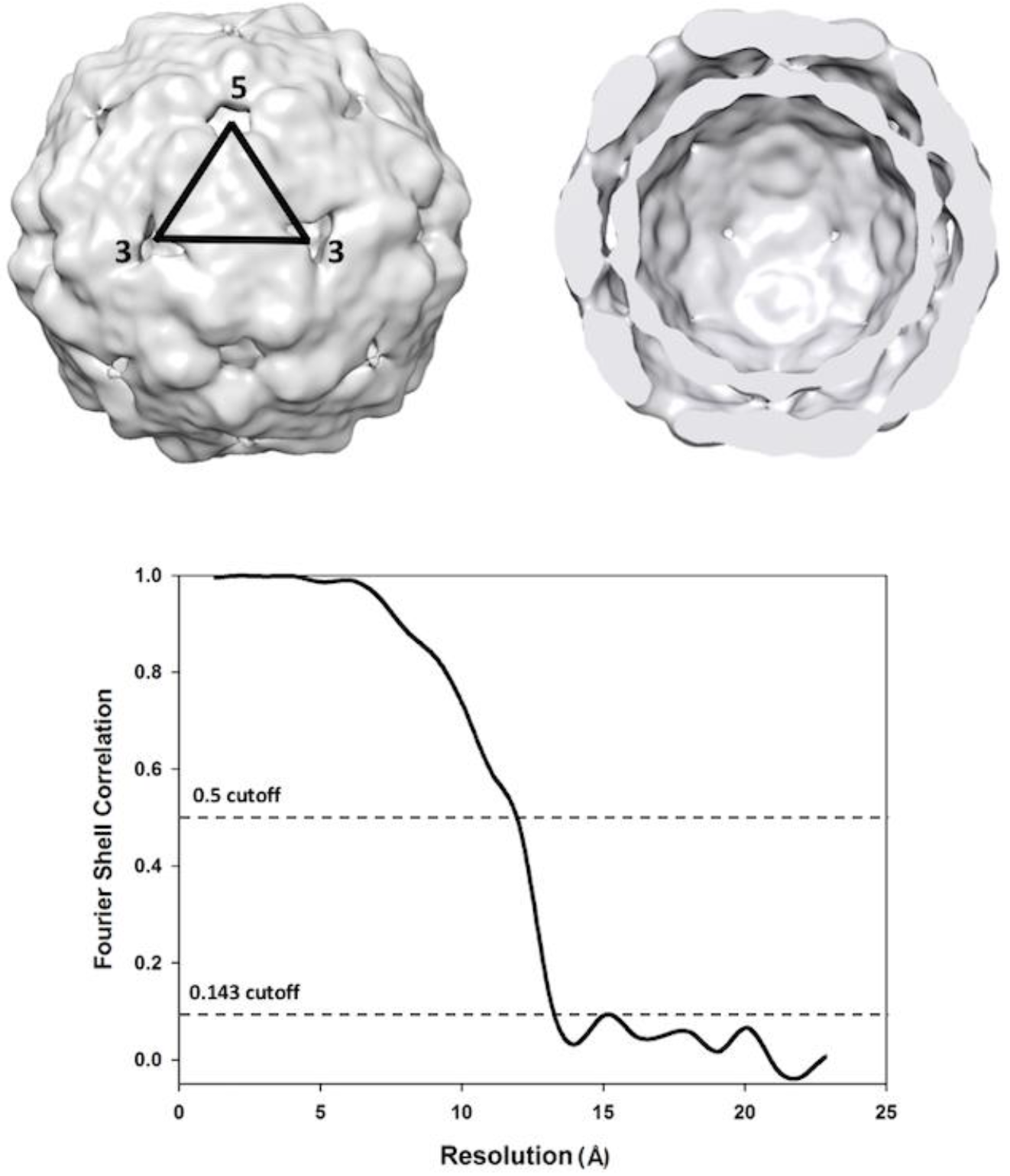
The cryo-EM structure of m2DVLP. The reconstructed cryo-EM map of the mD2VLPs was shown on the left. An asymmetric unit was indicated. On the right, the closest half of the density map has been removed to reveal the hollow structure. The resolution of the final reconstruction was determined to be 13.1Å using a gold standard resolution estimate at cutoff 0.143 (lower panel).

**Figure S6.**
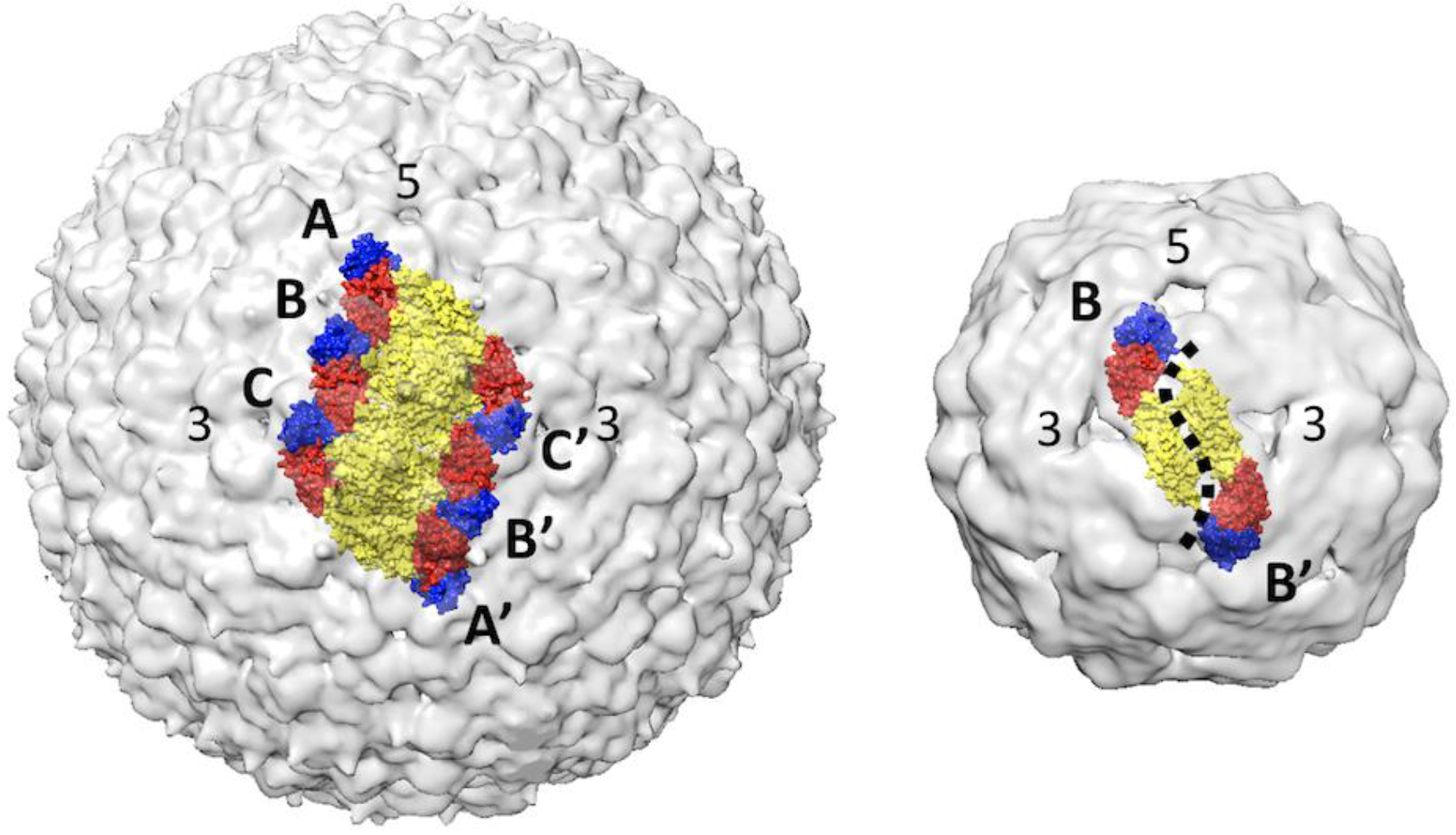
The E protein rafts in a virion (left) and mD2VLP (right). The E protein forming the rafts in the virion were shown in the left panel where the three individual E proteins in the asymmetric unit were labeled A, B, and C and the E proteins in the neighboring asymmetric unit are labeled A’, B’, and C’. The icosahedral 2-fold E protein dimers (B and B’) of mD2VLP moved apart from each other causing the groove (dashed line, right).

**Figure S7.**
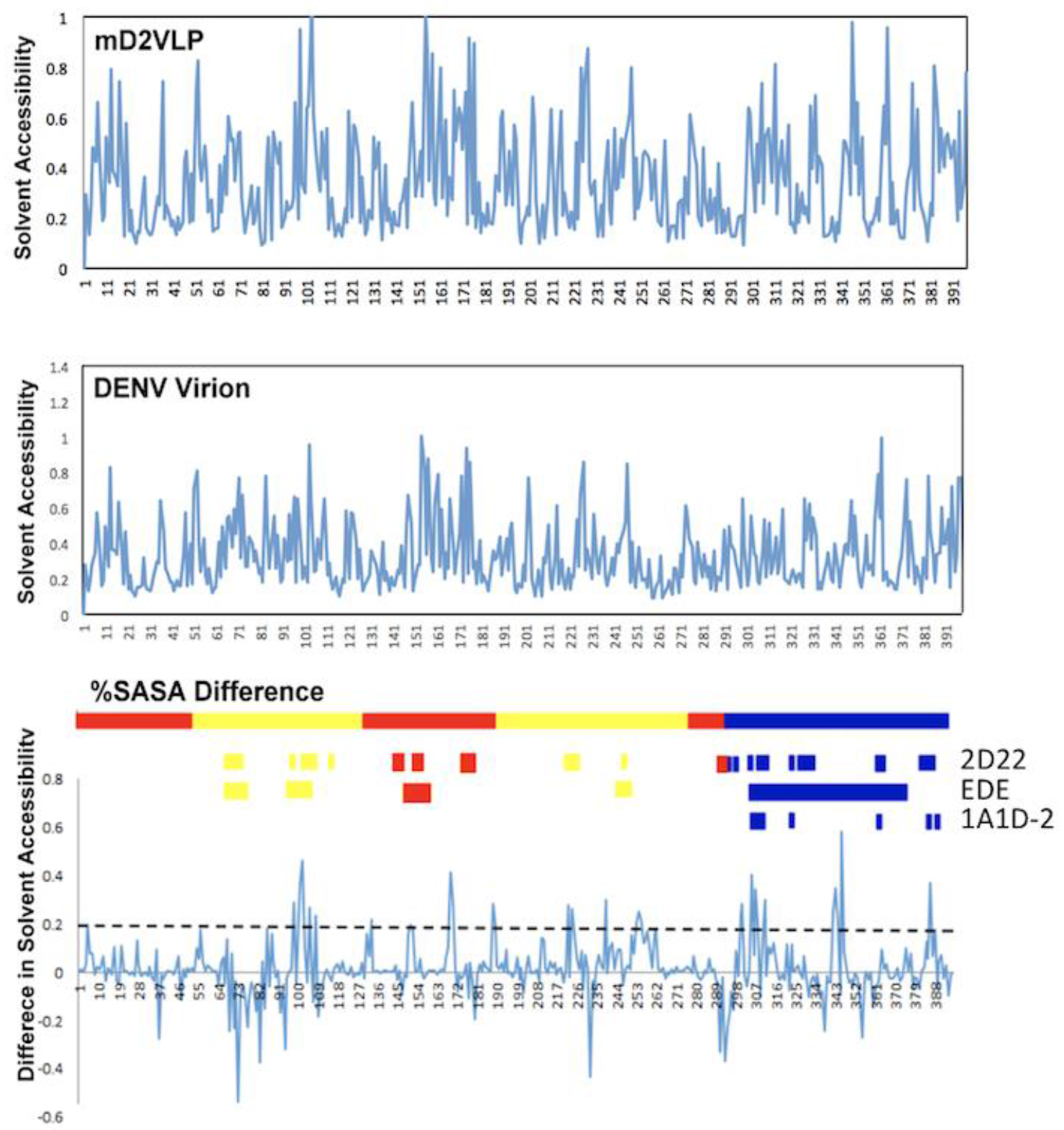
The solvent accessibility analyses. Solvent accessibility of the dengue particle soluble envelope (sE) protein amino acid (aa) residues 1-396 in DENV virion and mD2VLP (middle and top, respectively). The all-atom model of VLP was built using MODELLER^18^, and the all-atom model virion was built based on the cryo-EM structure of the mature dengue virus at 3.5Å resolution (PDB ID: 3J27). The relative solvent-accessible surface area (SASA) of both models was calculated using the POPS program. The plot shown represents the difference in relative solvent-accessible surface area (Δ%SASA) (Bottom). A positive value of %SASA means the residue becomes more exposed in mD2VLP than in DENV VLP. Domains I, II and III are highlighted in red, yellow and blue, respectively. The residues interacting with antibodies 2D22, EDE and 1A1D-2 are shown.

**Fig S8.**
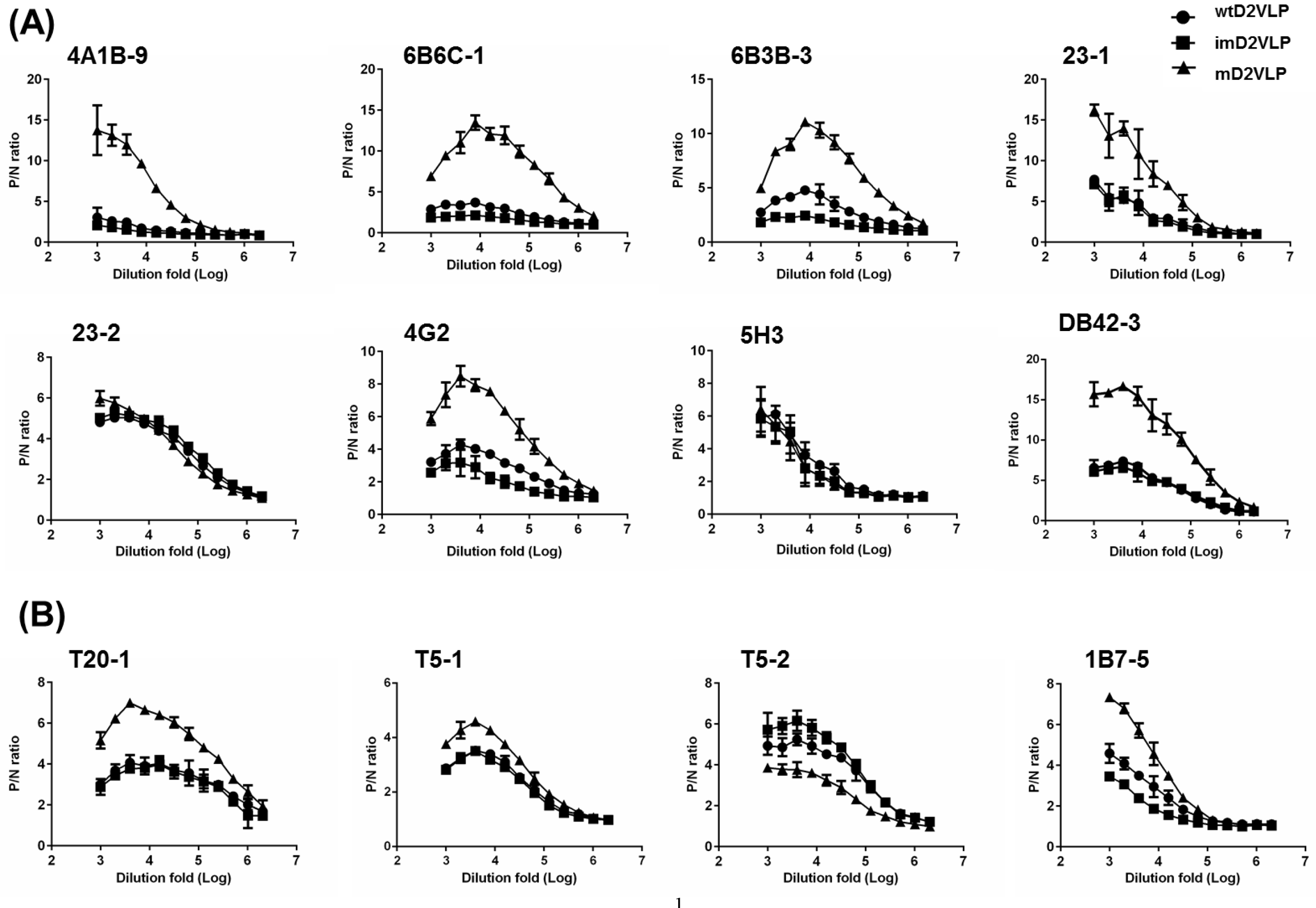
Binding activity curves of a panel of anti-E monoclonal antibodies (Mabs) to three types of D2VLP with different percentage of pr-peptide on particle surface. Binding curves for (A) eight group-reactive Mabs, (B) four subgroup-reactive Mabs, (C, D) four complex-or sub-complex-reactive Mabs, (E) four serotype-specific-reactive Mabs were determined by antigen-capture ELISA. Equal quantity of three types D2VLP were first titrated against purified D2VLP before adding to the wells. Mabs were serially 2-fold diluted starting from 1:1000 dilution. Data are expressed as P/N value by dividing the OD450 value from each dilution of Mab by the QD450 value from the control COS-1 culture supernatant. Data are means+SEM of duplicates from three representative experiments

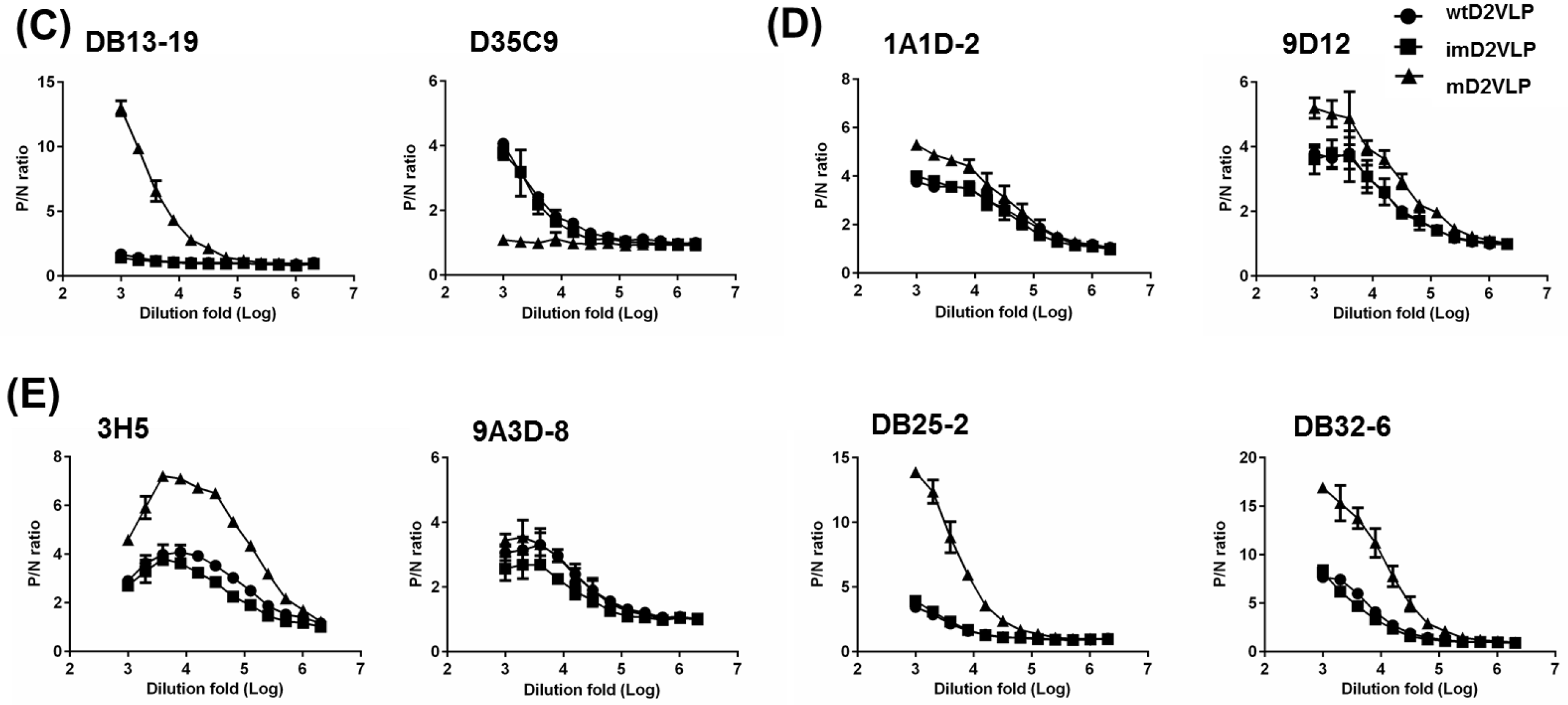

**Fig S9.**
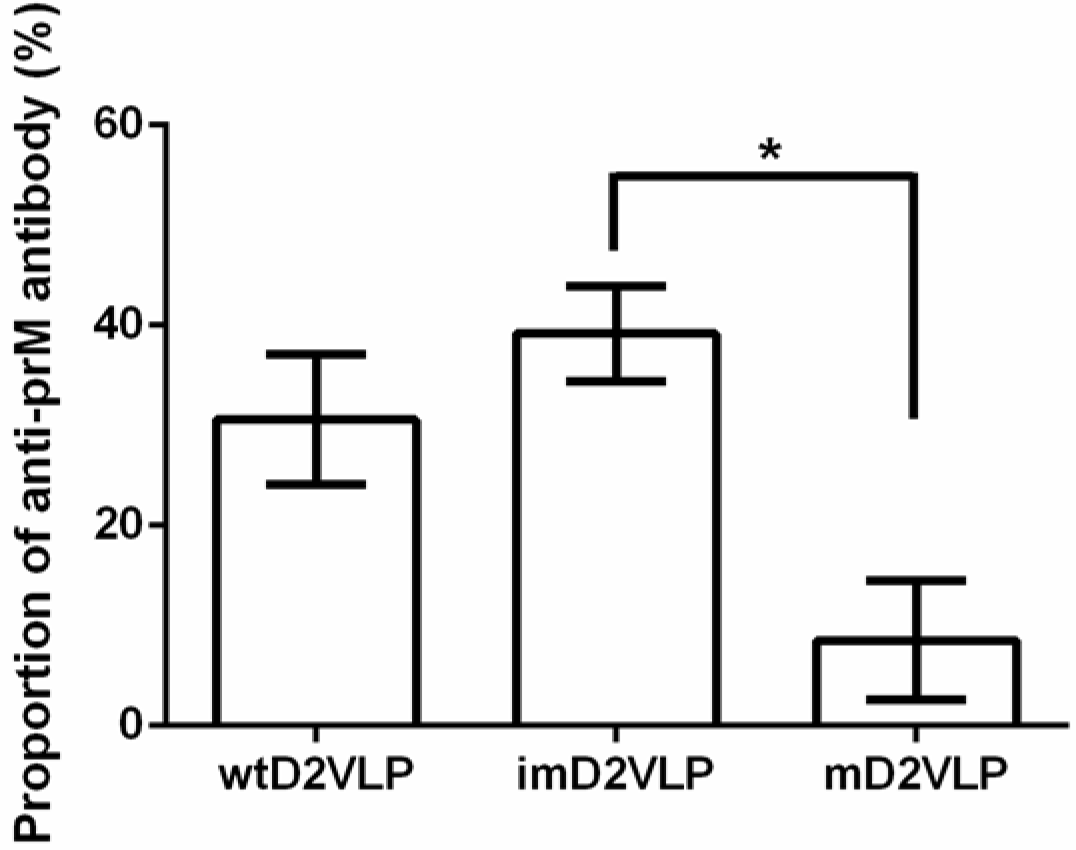
Proportions of anti-prM antibodies from both D2VLP immunization groups were measured using an epitope-blocking ELISA. Percent blocking of HRP-labeled anti-prM MAb (2H2-HRP) by sera from mice vaccinated with imD2VLP or mD2VLP was determined by the formula 100*[(GD450imD2VLP-QD450imD2VLP blocked by MAb 2H2)/OD450imD2VLP] using a 1:1000 dilution of mouse sera.

**Figure S10.**
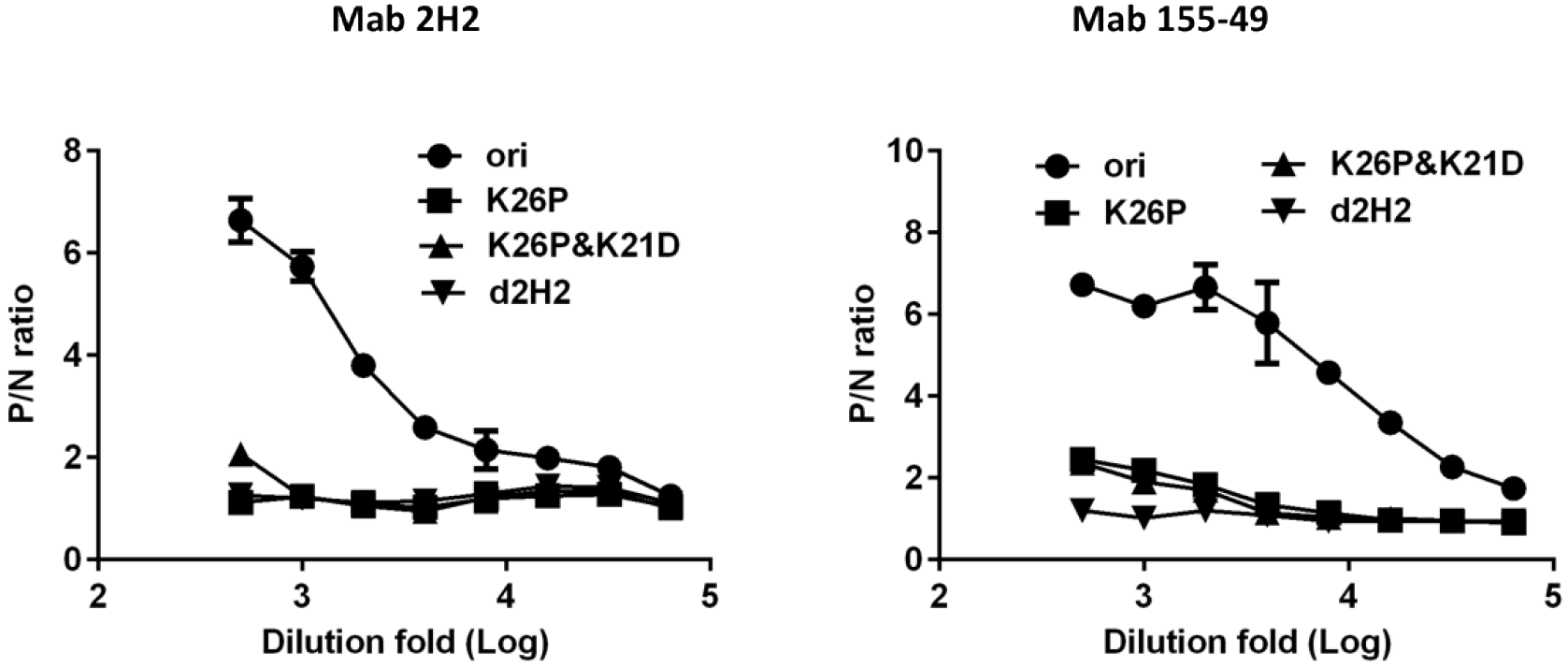
Epitope identification of MAbs 2H2 and 155-49. Single (K26P), double (K26P&K21D) and triple (K26P&K21D&F1A) mutations were performed on imD2VLP-expression plasmids by site-directed mutagenesis. Various D2VLP mutants were expressed in COS-1 cells by electroporation with plasmid DNAs as indicated and tissue culture media were clarified 3 days after transfection for antigen-capture ELISA. Substitutions of K26P on imD2VLP led to a significantly reduced binding activity of MAb 2H2; however, a weak binding can still be detected for MAb 155-49. Double mutation didn’t affect the binding of MAb 155-49 on imD2VLP, compared to a single mutation. Only the triple mutation completely abolished the binding of MAb 155-49 on imD2VLP. The antigens used were standardized at a single concentration with an optical density (OD) of 0.8, which was based on the standard curves generated by antigen-capture ELISA. Data are shown as means±SEM from three independent experiments. P/N ratio refers to the antibody binding magnitude between designated VLP-containing (P) and VLP-free culture supernatants (N) by dividing the absorbance of P by that of N.

**Fig S11.**
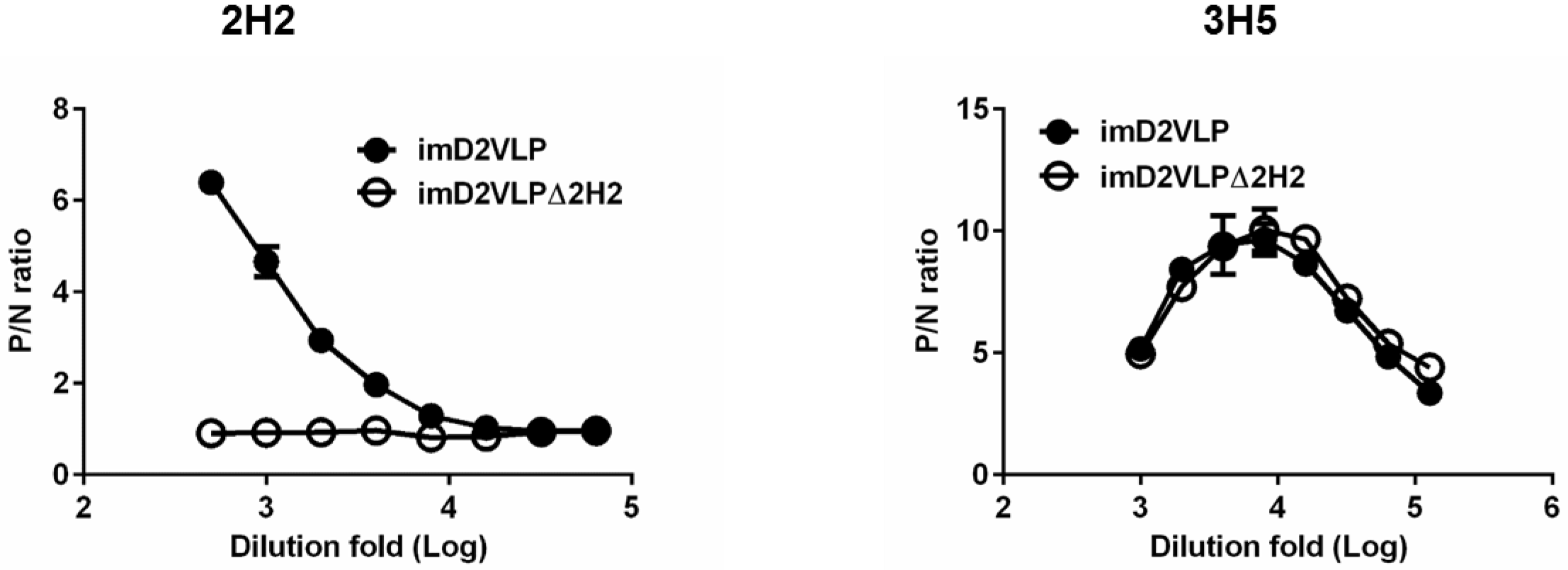
Serial dilutions of imD2VLP and mutant imD2VLP (Δ2H2), containing mutations at the MAb 2H2 binding site (F1A, K21D, K26P), were tested for binding with MAb 2H2 (left) and control antibody 3H5 (right) by ELISA. P/N ratios refer to the antibody binding magnitude between designated VLP-containing (P) and VLP-free culture supernatant (N) by dividing the absorbance of P by that of N.

**Figure S12.**
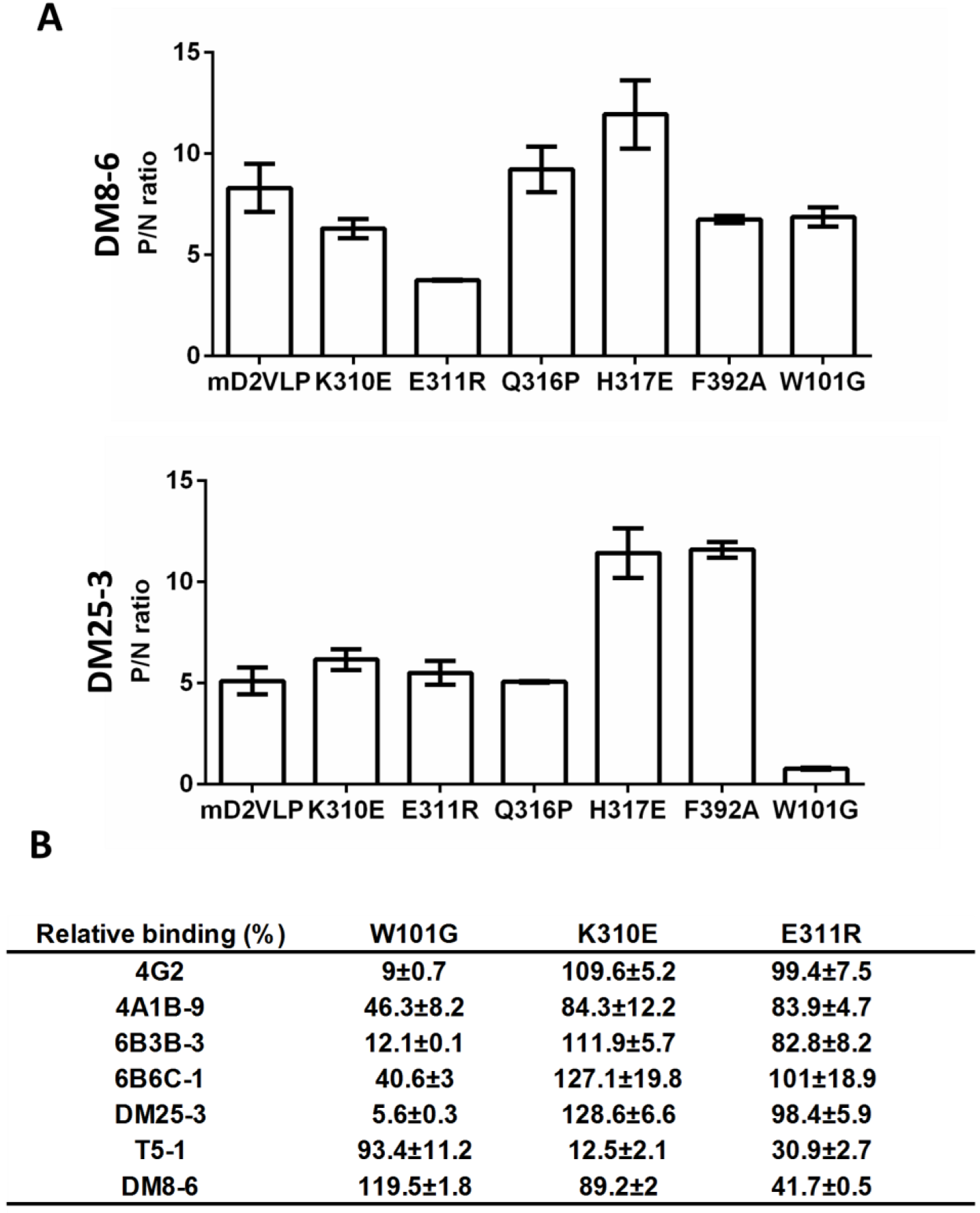
Identification of the neutralizing epitopes of MAbs DM25-3 and DM8-6. (A) Bar graph shows the decrease in MAb reactivity in a binding-ELISA for mD2VLPs with substitutions at the designated residues. Various mD2VLP mutants were expressed in COS-1 cells by electroporation with plasmid DNAs as indicated. Tissue culture media were harvested 3 days after transfection and clarified for antigen-capture ELISA. Substitutions at 1N103K, G104Q, K307E, E314R, T315H, P364R and W391G produced plasmids which did not secrete measurable VLP antigens into the tissue culture medium. Therefore, only mutant plasmids indicated secreted VLP antigens and were used for epitope mapping by binding ELISA. (Top) Substitutions of E311E led to a significantly reduced binding activity of neutralizing MAb DM8-6, compared to mD2VLP. (Bottom) Substitutions of W101G led to a complete loss of binding by MAb DM25-3. Data shown are a representative experiment of three independent experiments. (B) To ensure that the antigenic structures were intact, mutant VLPs including W101G, K310E and E311R were further confirmed by using a panel of DENV-2 MAbs, including group-cross-reactive antibodies (4G2, 4A1B-9, 6B3B-3, 6B6C-1, DM25-3) recognizing all four major pathogenic flavivirus serocomplexes; complex crossreactive antibodies (T5-1) recognizing all four DENV complex viruses, and type-specific antibody (DM8-6) recognizing DENV-2 only. The relative binding of MAbs was determined by dividing the 0D_450_ of mD2VLP mutants by that of mD2VLP. The data are presented as means ± SEM from three independent experiments with two replicates. The two-tailed Mann-Whitney *U* test was used to test statistical significance.

**Figure S13.**
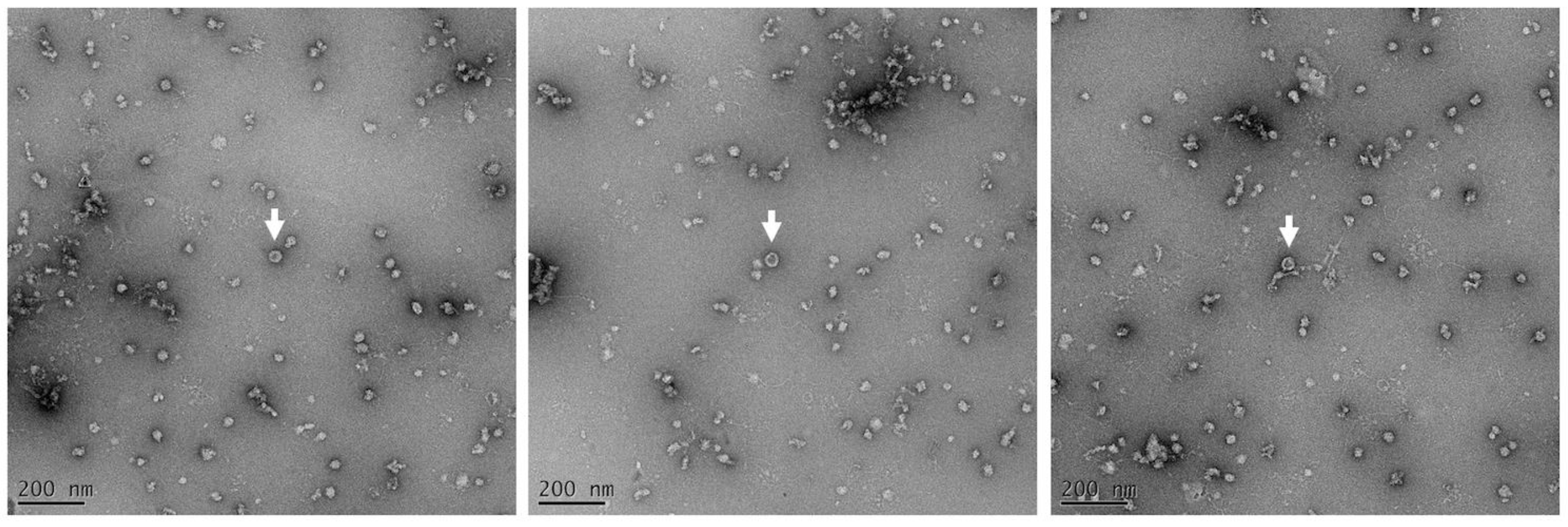
Transmission electron microscope (TEM) examination of negatively stained W101G-mutant mD2VLP. Purified W101G-mutant mD2VLP were adsorbed onto a glow-discharged carbon coated grid (EMS CF-200-Cu) for 1min, and inspected by TEM. The images were taken by a JEM1400 electron transmission microscope at a magnification of 30,000x using a 4k × 4k Gatan 895 CCD camera. The diameters of particles were between 30-32nm as measured by ImageJ software.

**Figure S14.**
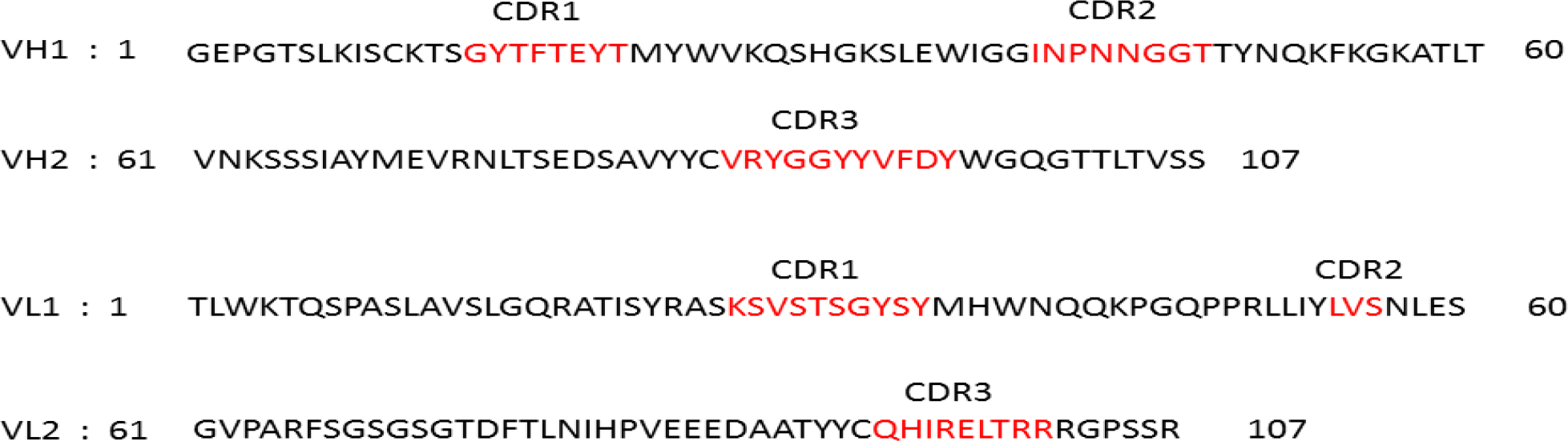
Amino acid sequences of variable regions of IgH and IgL of antibody DM25-3. The amino acids in the CDR regions are highlighted in red.

**Figure S15.**
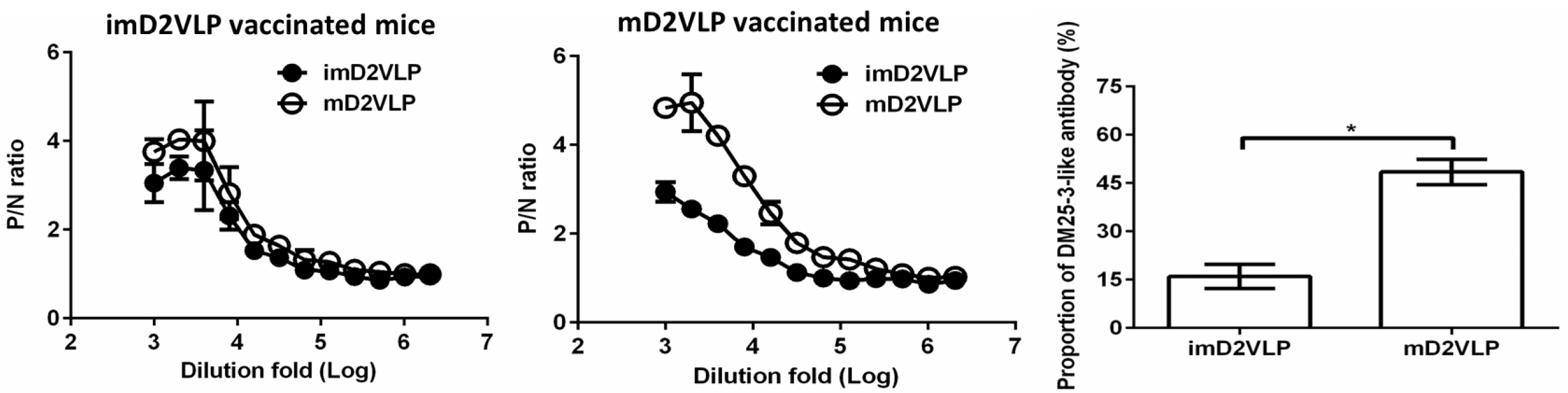
Binding ELISA was performed to test the reactivity of sera from mice immunized with two doses of imD2VLP or mD2VLP against equal amounts of imD2VLP and mD2VLP antigens. The difference in binding activity was converted to a bar chart at 1:1000-fold dilutions of mice sera. Proportions of DM 25-3-like antibodies from two different D2VLP immunization groups were calculated based on the formula 100x(OD_450_mD2VLP-OD_450_imD2VLP)/OD_450_mD2VLP. The data are presented as means ± SEM from three independent experiments with two replicates. The two-tailed Mann-Whitney *U* test was used to test statistical significance. *, *P* <0.05.

**Figure S16.**
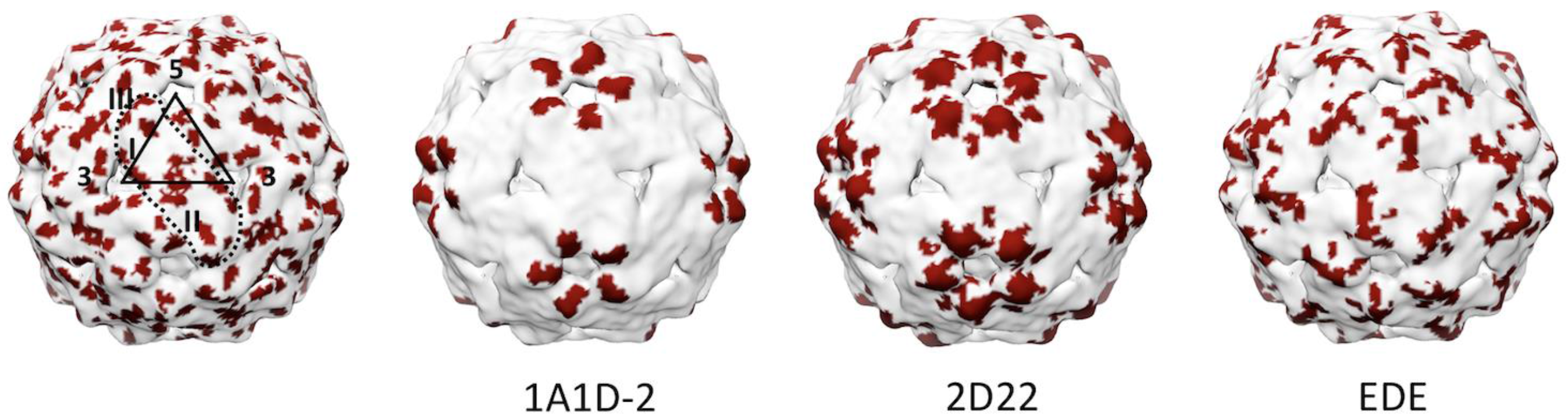
The neutralization-sensitive hotspot on mD2VLPs. (Left) The highly exposed residues (Δ%oSASAμ≥0.2) which were colored by dark red color are shown on the surface rendered VLP. The E dimer is outlined by a black dash line, the symmetry and the domains are labeled. The E epitopes interacting with MAb 1A1D-2 (center left), 2D22 (center right) and EDE (right) are colored in dark red on the surface showing that they were highly exposed on the mD2VLP surface and were highly overlapping with areas of highly exposed residues in mD2VLP

